# Mammalian mitochondrial mutational spectrum as a hallmark of cellular and organismal aging

**DOI:** 10.1101/589168

**Authors:** A. G. Mikhaylova, A. A. Mikhailova, K. Ushakova, E.O. Tretiakov, V. Shamansky, A. Yurchenko, M. Zazhytska, E. Zdobnov, V. Makeev, V. Yurov, M. Tanaka, I. Gostimskaya, Z. Fleischmann, S. Annis, M. Franco, K. Wasko, W.S Kunz, D.A. Knorre, I. Mazunin, S. Nikolaev, J. Fellay, A. Reymond, K. Khrapko, K. Gunbin, K. Popadin

## Abstract

Mutational spectrum of the mitochondrial genome (mtDNA) does not resemble signatures of any known mutagens and variation in mtDNA mutational spectra between different tissues and organisms is still incomprehensible. Since mitochondria is tightly involved in aerobic energy production, it is expected that mtDNA mutational spectra may be affected by the oxidative damage which is increasing with cellular and organismal aging. However, the well-documented mutational signature of the oxidative damage, G>T substitutions, is typical only for the nuclear genome while it is extremely rare and age-independent in mtDNA. Thus it is still unclear if there is a mitochondria - specific mutational signature of the oxidative damage. Here, reconstructing mtDNA mutational spectra for human cancers originated from 21 tissues with various cell turnover rate, human oocytes fertilized at different ages, and 424 mammalian species with variable generation length which is a proxy for oocyte age, we observed that the frequency of A_H_>G_H_ substitutions (_H_ - heavy chain notation) is positively correlated with cellular and organismal longevity. Moreover, this mutational bias from A_H_ to G_H_ affects nucleotide content at the fourfold degenerative synonymous positions leading to a deficit of A_H_ and excess of G_H_, which is especially pronounced in long-lived mammals. Taking into account additionally, that A_H_>G_H_ is sensitive to time being single stranded during mtDNA asynchronous replication and A>G is associated with oxidative damage of single-stranded DNA in recent bacterial experiments we propose that A_H_>G_H_ is a mutational signature of oxidative damage in mtDNA.

## Introduction

Various mutagens are characterized by their specific spectrum of the induced de novo mutations - mutational signatures. Investigation of these mutational signatures in the human nuclear genome helped to understand main cancer-specific mutagens (Alexandrov et al. 2020). Unlike the nuclear genome, key mutagens of the mitochondrial genome (mtDNA) are still unknown despite its 10-times increased mutation rate and numerous pathogenic mutations associated with diseases and aging (Wallace 2005).

Pan-cancer studies have shown that mitochondrial DNA (mtDNA) has a unique mutational spectrum which differs from all known mutational signatures in the nuclear genome and remains rather consistent across various cancer types (Yuan et al. 2017; Ju et al. 2014). For example, well known strong exogenous mutagens such as tobacco smoke in lung cancers or ultraviolet light in melanomas do not trigger significant damage in mtDNA, and thus the observed mitochondrial mutational spectrum has to be attributed to a potent unknown mutagen which can overwrite effects of others (Yuan et al. 2017; Ju et al. 2014). However, this potent mutagen is still unknown and it has not been shown if the mutagen is sensitive to internal metabolic cellular properties which are expected to vary among tissues and change with age. For example, G>T mutations, considered to be the hallmark of oxidative damage in nuclear DNA (COSMIC signature 18) (Fraga et al. 1990; Alexandrov et al. 2013; Kucab et al. 2019), do not significantly increase with age in mtDNA (Kennedy et al. 2013) and rather rare (Yuan et al. 2020), emphasising that mtDNA specific signature of oxidative damage, if exists, is still unknown. Interestingly, a similar gap of knowledge exists also on a comparative species level; although variation in mtDNA mutational signature between species has been shown in several works (Belle et al. 2005; Montooth and Rand 2008), no common explanation has been proposed yet.

It has been recently shown that, because of high mtDNA mutation rate and high mtDNA copy number per cell, somatic variants in mtDNA can provide crucial information allowing us to trace different cellular lineages in our body (Ludwig et al. 2019). Here, taking into account potential association of the mtDNA endogenous mutagens with cellular properties, we hypothesize further that the relative frequency of different mtDNA substitutions (i.e. mtDNA mutational spectrum) may contain a functional signature of cellular properties related to the level of metabolism and aging. Indeed, mitochondria are the organelles determining the cell-specific level of oxidative metabolism and, as a consequence, associated mutagens are expected to directly and strongly affect mtDNA (Ericson et al. 2012).

Here, focusing on cellular longevity (Tomasetti and Vogelstein 2015; Tomasetti et al. 2017) we observed differences in mtDNA mutational spectrum between tissues with short- and long-lived cells: frequency of A_H_>G_H_ substitutions positively correlates with cellular longevity. Next, we extended our analyses to human mother-offspring pairs and other mammalian species. As the mtDNA of all mammals is inherited exclusively through the oocytes, and oocyte division is stopped after birth, the species-specific generation length is close enough to the longevity of oocytes from different mammalian species (i.e. the time the mtDNA spent in the dormant oocyte) and could therefore be correlated with the specific mtDNA signature of longevity. Indeed, we observed that the frequency of A_H_>G_H_ substitutions positively correlates with organismal longevity. Finally, the discovered mutational spectrum left fingerprints on the neutral nucleotide content of the whole mitochondrial genomes as an excess of G_H_ and a deficit of A_H_ which is more pronounced in long-versus short-lived mammals. Altogether, using a collection of somatic mtDNA mutations from human cancers, de novo germline mutations from the mother-offspring pairs, polymorphic synonymous substitutions from hundreds of mammalian species, as well as nucleotide content in whole mitochondrial genomes of mammals, we observed one universal trend: the frequency of A_H_>G_H_ substitutions positively correlates with cellular and organismal longevity. We propose that A_H_>G_H_ is a mtDNA specific mutational signature of oxidative damage, which is increasing with cellular and organismal aging.

## Results

### (1) mtDNA mutational spectrum in human cancers changes during tumorigenesis and is associated with cell turnover rate in ancestral tissues

Results of recent comprehensive survey of somatic mtDNA mutations in human cancers demonstrated that mtDNA mutational spectra are conserved across different cancer types with rare transversions and clear prevalence of C_H_>T_H_ and A_H_>G_H_ transitions (Yuan et al. 2020) (hereafter _H_ marks heavy strand notation). However, minor variations in mutational spectra might be indicative of cancer-stage or tissue-specific metabolic rates. To test whether mtDNA mutational spectrum changes during cancerogenesis we used a collection of 7611 somatic mitochondrial mutations from 37 cancer types grouped into 21 ancestral tissues (Yuan et al. 2020). To rank all somatic mutations from early to late, we evaluated the variant allele frequency (VAF) of each variant: high VAFs being a signature of early mutations (originated early during cancerogenesis and thus presented in the majority of cancer cells) and low VAFs being a signature of late mutations (originated at the last stages of cancerogenesis and thus presented in a subclone of the cancer cells) (Supplemental Mat 1.1, Figure S1A). Splitting all variants by the median VAF (4.5%), we observed a significant decrease in transition/transversion ratio (hereafter Ts/Tv) among late variants (from 14.9 to 10.3, Supplemental Mat 1.1, Figure S1B). This decrease in Ts/Tv among late variants is observed not only in the total set of 7611 somatic mutations but also within the majority of cancer subtypes and even within individual samples (Supplemental Mat 1.1, Figure S1B). Transversions are more common among late variants and, despite numerous quality control steps (Yuan et al. 2020), they might be enriched in false-positive variants, partially driving our observation. To avoid a contribution of the potential false-positive transversions, we next focused on two of the most common transitions: C_H_>T_H_ and A_H_>G_H_ and compared their relative time of origin (i.e. VAFs) in individual samples. We observed that A_H_>G_H_ occurs slightly earlier (i.e. has higher VAF) as compared to C_H_>T_H_ (Supplemental Mat 1.1), providing an independent line of evidence for mutational changes associated with tumorigenesis. Thus, mtDNA mutational spectrum is changing during tumorigenesis: both Ts/Tv and A_H_>G_H_ / C_H_>T_H_ are decreasing from early to late variants suggesting that mitochondrial mutagenic microenvironment is dynamic throughout tumorigenesis.

Many somatic mitochondrial mutations, annotated in human cancers, would have occurred before the tumorigenesis (Supplemental Mat 1.1, Figure S1A) and thus can be indicative of healthy tissue properties. One of the most prominent tissue-specific properties, linked to both mutagenesis (Tomasetti and Vogelstein 2015) and mitochondrial metabolism, is the turnover rate of stem cells in a given tissue. Based on extensive literature search we derived an approximate turnover rate of stem cells in each of 21 normal tissue samples, ancestral to the analyzed cancer types (Supplemental Mat 1.2, Table S1). Correspondingly, we split all these tissues, which are representative of different cancers, into three categories:

fast-replicating - with stem cell turnover rate less than a month (uterus, colon/rectum, stomach, cervix, esophagus, head/neck, lymphoid and myeloid tissues); intermediate - with stem cell turnover rate higher than a month and less than ten years (breast, prostate, skin, bladder, biliary, pancreas, liver and kidney) and slow-replicating - with stem cell turnover rate higher than ten years (thyroid, lung, bone/soft tissue, central nervous system (CNS) and ovary). Estimating the mutational spectrum for each category we observed an increase in Ts/Tv ratio from 10.0 in a fastly-dividing category, to 12.6 in intermediate, and to 14.5 in the slow-dividing one (Supplemental Mat 1.2, Figure S1C, left panel). The observed difference is robust to jackknife resampling and is maintained mainly by a common transition A_H_>G_H_ (Supplemental Mat 1.2, Figure S1C, middle panel). Comparing the frequencies of both these transitions with each other we observed an increase in the ratio A_H_>G_H_ / C_H_>T_H_ from fast-to slow-replicating tissues (A_H_>G_H_ / C_H_>T_H_: 0.56, 0.58, 0.61 for fast-, intermediate and slow-replicating tissues; Supplemental Mat 1.2, Figure S1C, right panel). Thus, Ts/Tv is increasing from fast-to slow-replicating tissues and this effect is associated with an increased fraction of A_H_>G_H_ transitions.

Putting together the two observations described above: the increased A_H_>G_H_ in early stages of cancer (Supplemental Mat 1.1) and increased A_H_>G_H_ in slow dividing tissues (Supplemental Mat 1.2), we expect that a probability of an A_H_>G_H_ transition is a function of both (i) relative time of origin of each somatic mutation, approximated as 1 minus VAF, and (ii) tissue-specific turnover rate. To test it, we performed multiple logistic regression and observed that the probability of a A_H_>G_H_ transition indeed decreases with both time of cancerogenesis and turnover rate of the tissue (Figure 1, Supplemental Mat 1.3, Table S2). The observed result was robust to an introduction of the factorial variable encoding fast-replicating cancers, to an elimination of 25% of the rarest potentially false positive variants, to a rerunning of this analysis only with transitions and to an addition to the model the patient age and the number of mitochondrial copies per cancer samples (Supplemental Mat 1.3, Table S2). Thus, A_H_>G_H_ substitutions are more frequent (i) at earlier stages of tumorigenesis and (ii) in cancers derived from slow-replicating tissues.

**FIGURE 1:**
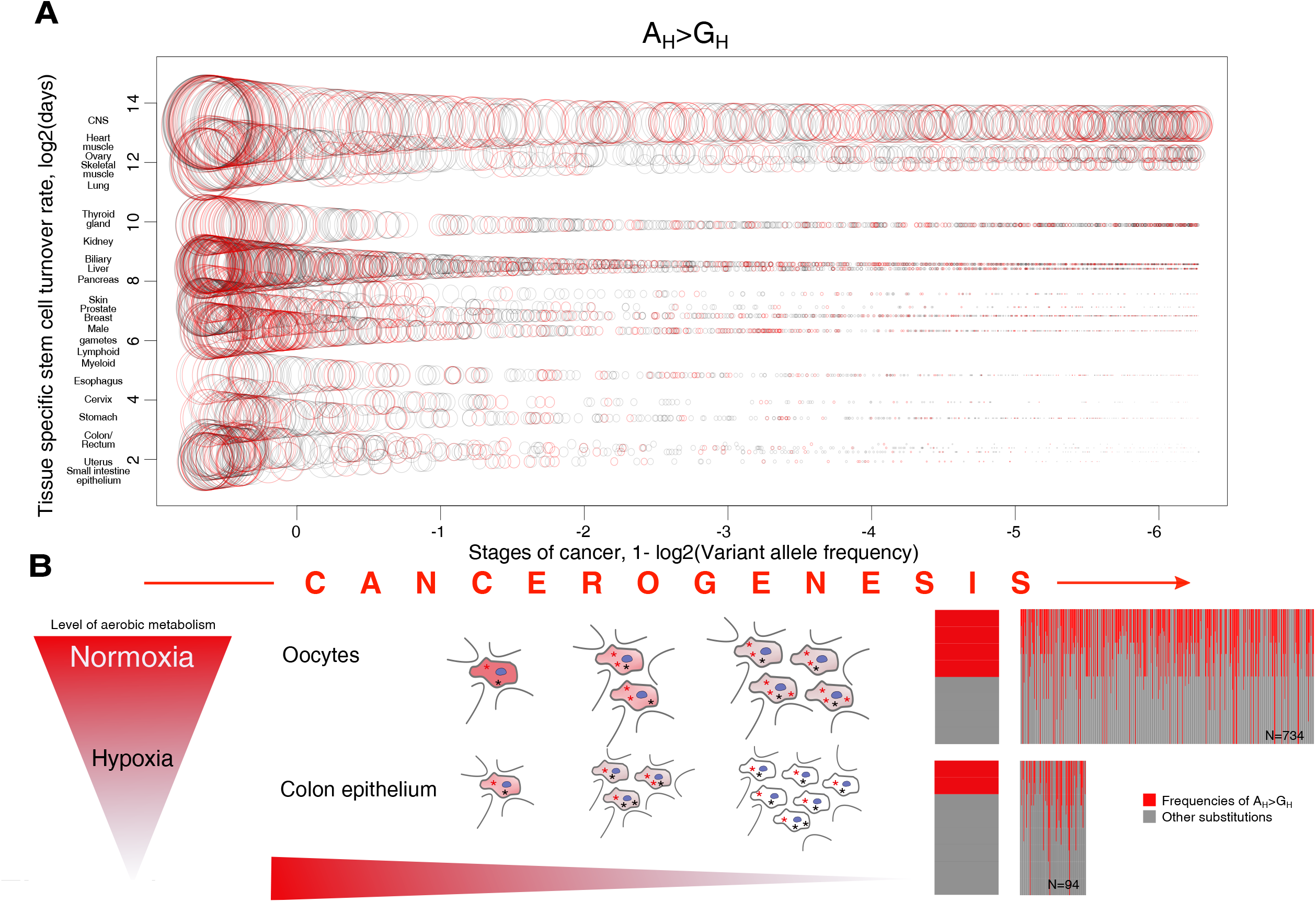
mtDNA mutational spectra is changing between tissues and during carcinogenesis. 1A. The probability of A_H_>G_H_ is higher in early stages of cancer (when cells are more normoxic and slow dividing) and in tissues with low turnover rate. X axis reflects the stages of cancerogenesis approximated by 1 - VAF. Y axis marks the stem cell turnover rate. Each circle reflects one mutation. The size of the circles is proportional to a probability that a given mutation is A_H_>G_H_, where the probability is obtained from the logistic regression model 1A (Supplemental Mat 1.3) and normalized to 0-1 range. Red circles mark A_H_>G_H_, grey circles - all other substitutions. It is seen that circles (probability of A_H_>G_H_) are decreasing from left to right reflecting increased hypoxia in more advanced stages of cancers and from top to bottom, reflecting a gradient of tissues with low to high turnover rate. 1B. The probability of A_H_>G_H_ is decreased in hypoxic samples. The scheme on the left demonstrates expected changes in normoxia level among different tissues (for example slow dividing oocytes and fastly dividing colon epithelium) and stages of cancer (all cancers slowly become more hypoxic with time). A fraction of A_H_>G_H_ (red asterisks) reflects oxidative damage at a given moment in cancer development: from the highly aerobic, normoxic (red background color) early stages to late, hypoxic stages (grey background color). The barplots on the right demonstrate differences in the frequencies of A_H_>G_H_ between normoxic samples (734 samples with Buffa score lower than 32) and hypoxic samples (94 samples with Buffa score higher or equal to 32). Each bar corresponds to one sample where the red part reflects the sample-specific frequency of A_H_>G_H_.

The changes in mutational spectra during the tumorigenesis as well as between slowly and fastly dividing tissues might be associated with different levels of aerobic metabolism, ranging from hypoxia to normoxia. It has been shown, for example, that mtDNA in hypoxic colon cancer exhibited threefold decreased mutation rate as compared to normoxic adjacent non-tumor tissue (Ericson et al. 2012). This effect was mainly driven by the decrease in both common transitions (C_H_>T_H_ and A_H_>G_H_), which are expected to be associated with oxidative damage and thus become rarer in hypoxic cancer conditions. Taking into account that practically all cancer types are becoming more hypoxic with time (Rosario et al. 2018) we interpret the fraction of A_H_>G_H_ as a marker of oxidative damage at a given moment in cancer development: from the highly aerobic, normoxic, early stages (high fraction of A_H_>G_H_) to late, hypoxic stages (low fraction of A_H_>G_H_). Using hypoxia scores derived for many individual cancer samples in the framework of the ICGC/TCGA Pan-Cancer Analysis of Whole Genomes (PCAWG) Consortium (Bhandari et al. 2020) we were able to test directly an association between hypoxia and fraction of A_H_>G_H_. Analysis of the whole dataset showed only suggestive evidence of correlation (Spearman’s rho = -0.046, p-value = 0.1869, N = 828), however, this relationship became more evident when we compared the upper decile of the most hypoxic cancers (frequency of A_H_>G_H_ is 0.20, N = 94 with Buffa score higher or equal to 32) versus all other, less hypoxic cancers (frequency of A_H_>G_H_ is 0.28, N = 734, Buffa score lower than 32) (p = 0.0047, Mann-Whitney U-test, Figure 1B right panel). Altogether, we suggest that mtDNA mutational spectrum is sensitive to the level of the aerobic metabolism by means of A_H_>G_H_ transitions susceptible to an oxidative damage, which is higher in early-stage cancers and in slow-dividing tissues.

### (2) mtDNA mutational spectrum changes with female reproductive age

Analyzing cancer data we observed that mtDNA mutational spectrum, and particularly a high fraction of A_H_>G_H_ substitutions, marks tissues with slow turn-over rate (Figure 1A). We suggest that high fraction of

A_H_>G_H_ can be a result of increased oxidative damage in slow-dividing long-lived cells (Figure 1B). If so, we can expand our logic and switch from somatic tissues to human oocytes, which demonstrate a high degree of variation in longevity (time of fertilization) and may have an altered mtDNA mutational spectrum.

By comparing mitochondrial variants in offsprings and mothers, it is possible to reconstruct the de novo mitochondrial mutations that occurred in oocytes. It has been shown that the number of de novo mtDNA mutations in children is increasing with maternal age (Rebolledo-Jaramillo et al. 2014; Wei et al. 2019; Zaidi et al. 2019), that might be attributed to oocyte aging (oocyte longevity). However, no age-related changes in mtDNA mutational signatures have been discovered yet. In order to test potential switch in mtDNA mutational spectra that occurs when oocyte age, we reanalyzed all of the de novo germline mutations from three recent studies of mother-offspring pairs (Rebolledo-Jaramillo et al. 2014; Wei et al. 2019; Zaidi et al. 2019). All the analyses of de novo mutations in mother-offspring pairs were only suggestive due to low available sample sizes but consistently showed a trend of A_H_>G_H_ rate increasing with oocyte longevity. Having access to the information about mothers’ age at the moment of giving birth from studies of Rebolledo-Jaramillo and Zaidi (Rebolledo-Jaramillo et al. 2014; Wei et al. 2019; Zaidi et al. 2019) we observed that A_H_>G_H_ de novo mutations were characterized by the highest age of reproduction (Supplemental Mat 2). Having access to the VAF values of de novo mtDNA mutations from the study of Wei et al 2020 (Wei et al. 2019) we observed the later onset of A_H_>G_H_ (i.e. lower VAFs of A_H_>G_H_) as compared to other mutations (Supplemental Mat 2). Despite the low sample size all three studies are in line with our hypothesis that a high fraction of A_H_>G_H_ marks aged, long-lived oocytes.

### (3) Mutational spectrum, derived from the intra-species mtDNA polymorphisms of mammals, is shaped by generation time

Until now we have been comparing different cells and tissues from the human body and observed that the fraction of mtDNA A_H_>G_H_ transitions out of twelve possible single-nucleotide substitutions positively correlates with cell longevity. However, if cell longevity is the main driver of the observed association, we may extend our logic further and analyze oocytes from different mammalian species. Oocytes are the only lineage through which mtDNA is transmitted (with rare exceptions of paternal inheritance) from generation to generation in mammals (Sato and Sato 2017). Mammalian oocytes are arrested from birth to puberty, which takes weeks (mouse) or decades (humans) (Von Stetina and Orr-Weaver 2011) and allows us to use the species-specific generation length as a good proxy for oocyte longevity in different mammalian species. Mitochondrial polymorphisms in mammals are especially advantageous for the reconstruction of species-specific mtDNA mutational spectrum because they were extensively sequenced in numerous ecological, evolutionary, and population genetics studies. As an example, the *MT-CO1* gene was selected for the DNA barcoding project (Hebert et al. 2003). Several studies revealed that there is some variation in mtDNA mutational spectra between different species (Montooth and Rand 2008; Belle et al. 2005), however, no overarching explanation has been suggested so far.

For 611 mammalian species, we collected all available mtDNA polymorphisms and reconstructed normalized 12-component mutational spectra as a probability of each nucleotide to mutate to any other nucleotide in the most neutral - fourfold degenerative synonymous sites (see Methods and Supplemental Mat 3.1, 3.2). The average mutational spectrum for all mammalian species (Figure 2A) demonstrates an excess of C_H_>T_H_ and A_H_>G_H_ substitutions which is similar to previous studies (Ju et al. 2014; Yuan et al. 2020).

**FIGURE 2:**
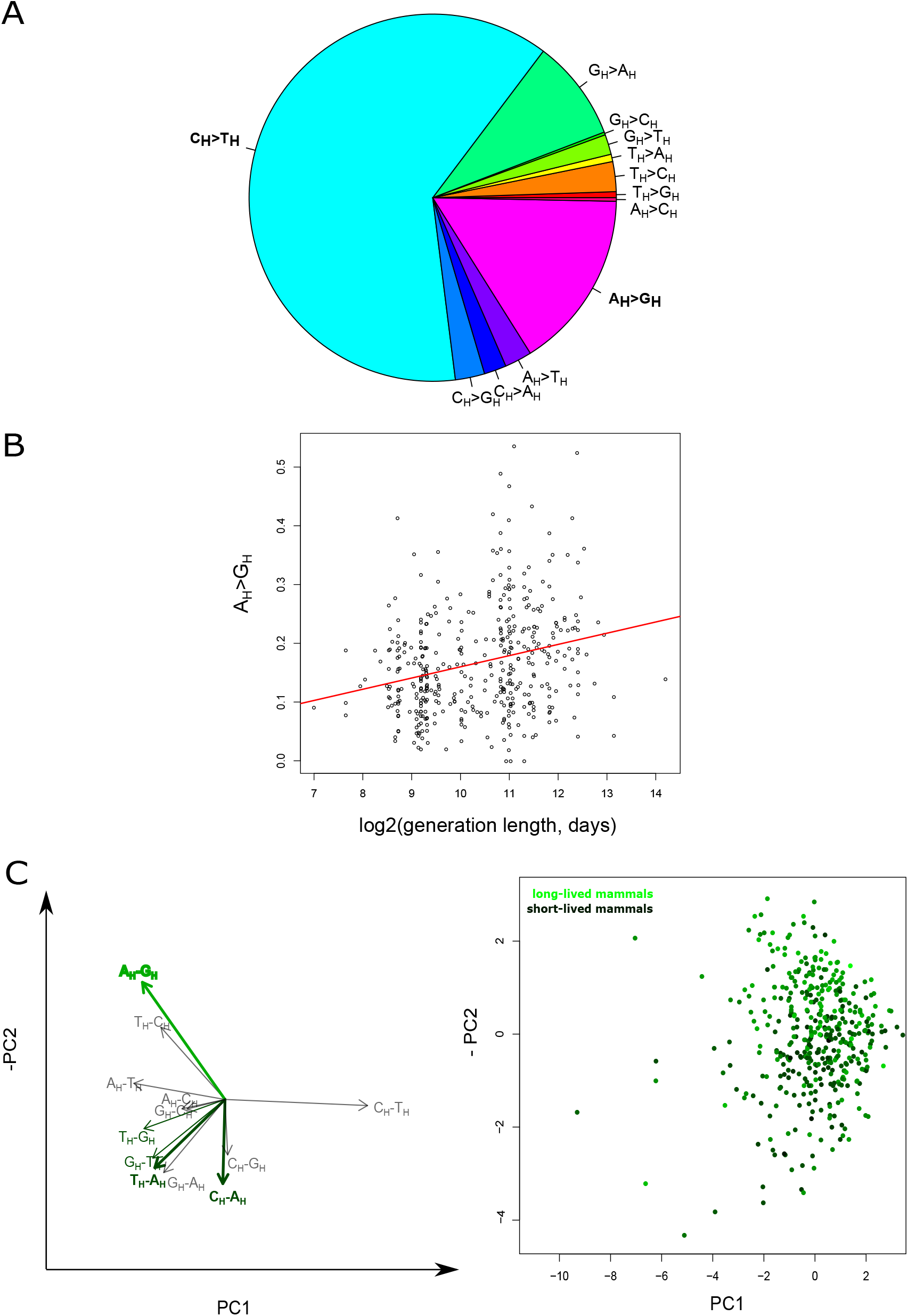
variation in mammalian mutational spectrum is driven by generation length. 2A. Average mtDNA mutational spectrum of mammalian species (N = 611). Mutational spectrum as a probability of each nucleotide to mutate to each other based on the observed and normalized frequencies of twelve types of nucleotide substitutions in four fold degenerate synonymous sites of all available within-species polymorphisms of mtDNA protein-coding genes. 2B. Mutational spectra vary with species-specific generation length (N = 424). Frequency of A_H_>G_H_ is the best type of substitutions, correlated with generation length. 2C. The principal component analysis of mtDNA mutational spectra of mammalian species (N = 424). Left panel: the biplot of the principal component analyses (first and the second components explains 16% and 12% of variation correspondingly). C_H_>T_H_ has the highest loading on the first principal component while A_H_>G_H_ has the highest loading on the second principal component. Note that we reverted PC2 to make it positively correlated with generation length. Right panel: The second principal component correlates with generation length in mammals. Generation length is color-coded from dark green (the shortest generation length) to light green (the longest generation length).

To uncover the nature of mitochondrial mutagenesis shaping the species-specific variation in mutational spectra we focused on *MT-CYB* gene, which contained 56% of all analyzed mutations (39,112 out of 70,053 used to draw Figure 2A) and compared the *MT-CYB*-derived spectrum between species with variable life-history traits. We used the well-characterized mammalian generation length as a metric (Pacifici et al. 2013; Tacutu et al. 2013) which is associated with oocyte longevity and also with numerous ecological (body mass, litter size, effective population size) and physiological (basal metabolic rate) parameters of mammalian species (Ollason 1987; Damuth 1987). As the first and simplest approximation of the mutational spectrum, we used Ts/Tv.

For 424 mammalian species with known Ts/Tv and generation length, we observed a positive correlation between those two parameters (Supplemental Mat 3.3). To test the robustness of this correlation we performed several additional analyses and showed that the results were not affected by (i) splitting of the mammalian species into groups by generation length and families, (ii) phylogenetic inertia, (iii) the total number of reconstructed mutations used to estimate the species-specific mutational spectrum (Supplemental Mat 3.3).

To understand which substitution types shaped the observed correlation between Ts/Tv and generation length we performed twelve pairwise rank correlation analyses. We observed that only A_H_>G_H_ frequency positively correlated with the generation length (Spearman’s rho = 0.252, p value = 1.188e-07) (Figure 2B), while several rare transversions showed a negative correlation (T_H_>A_H_, T_H_>G_H_, C_H_>A_H_ and G_H_>T_H_: all Spearman’s rhos < -0.17, all p-values are < 0.0003). Inclusion of all five types of substitutions into a multiple linear model showed A_H_>G_H_ being the strongest component associated with generation length; the result was also robust to phylogenetic inertia (Supplemental Mat 3.3).

In order to derive mtDNA mutational signatures in an unsupervised way we performed principal component analysis (PCA) of mutational spectra of 424 mammalian species. We observed that the first component is mainly driven by C_H_>T_H_ substitutions, whereas the second is driven mainly by A_H_>G_H_ substitutions (Figure 2C left panel). We assessed if the first ten principal components were correlated with generation length and observed that only the second component was significantly correlated (Figure 2C right panel).

Interestingly, the correlation of the second principal component with generation length was higher as compared to the sole effect of A_H_>G_H_ frequency, suggesting that the second component could reflect a complex nature of longevity-associated mutational signature (Supplemental Mat 3.5). Additional analyses which take into account different ways to normalize the frequency of A_H_>G_H_: by the most common transition C_H_>T_H_ (Supplemental Mat 3.6) or by the all types of substitutions with ancestral nucleotide being A_H_ (Supplemental Mat 3.7) confirmed the robustness of our results.

Altogether, our analyses of within-species neutral polymorphisms in hundreds of mammalian species demonstrated a clear sensitivity of mtDNA mutational spectrum, i.e. the frequency of A_H_>G_H_ substitutions, to the generation length.

### (4)The long-term mutational bias affects nucleotide composition of complete mammalian mitochondrial genomes

Mutational bias, if stronger than selection and if stable for a long period of time so that species are close to mutation-selection equilibrium, is expected to change genome-wide nucleotide content. We expect that increased frequency of A_H_>G_H_ (Figure 2) would decrease frequencies of A_H_ and increase frequencies of G_H_ in effectively neutral positions of long-lived species. Using 650 complete mitochondrial genomes of mammalian species we estimated their nucleotide content in the most neutral - synonymous fourfold degenerate - positions: in 13 protein-coding genes without overlapped regions. Testing all four pairwise correlations between the species-specific generation length and the nucleotide content we observed two strongest correlations: negative with A_H_ and positive with G_H_ (Figure 3A; Supplemental Mat 4.1, Table S3). Inclusion of all four types of nucleotide frequencies into the multiple model confirmed the importance of A_H_ and G_H_ only, the effect of which was also robust to the phylogenetic inertia (Supplemental Mat 4.1). Thus we concluded that mtDNA of long-versus short-lived mammals is more A_H_ poor and G_H_ rich (Figure 3A), which is in line with the more intensive A_H_>G_H_ mutagenesis in the former (Figure 2).

**FIGURE 3:**
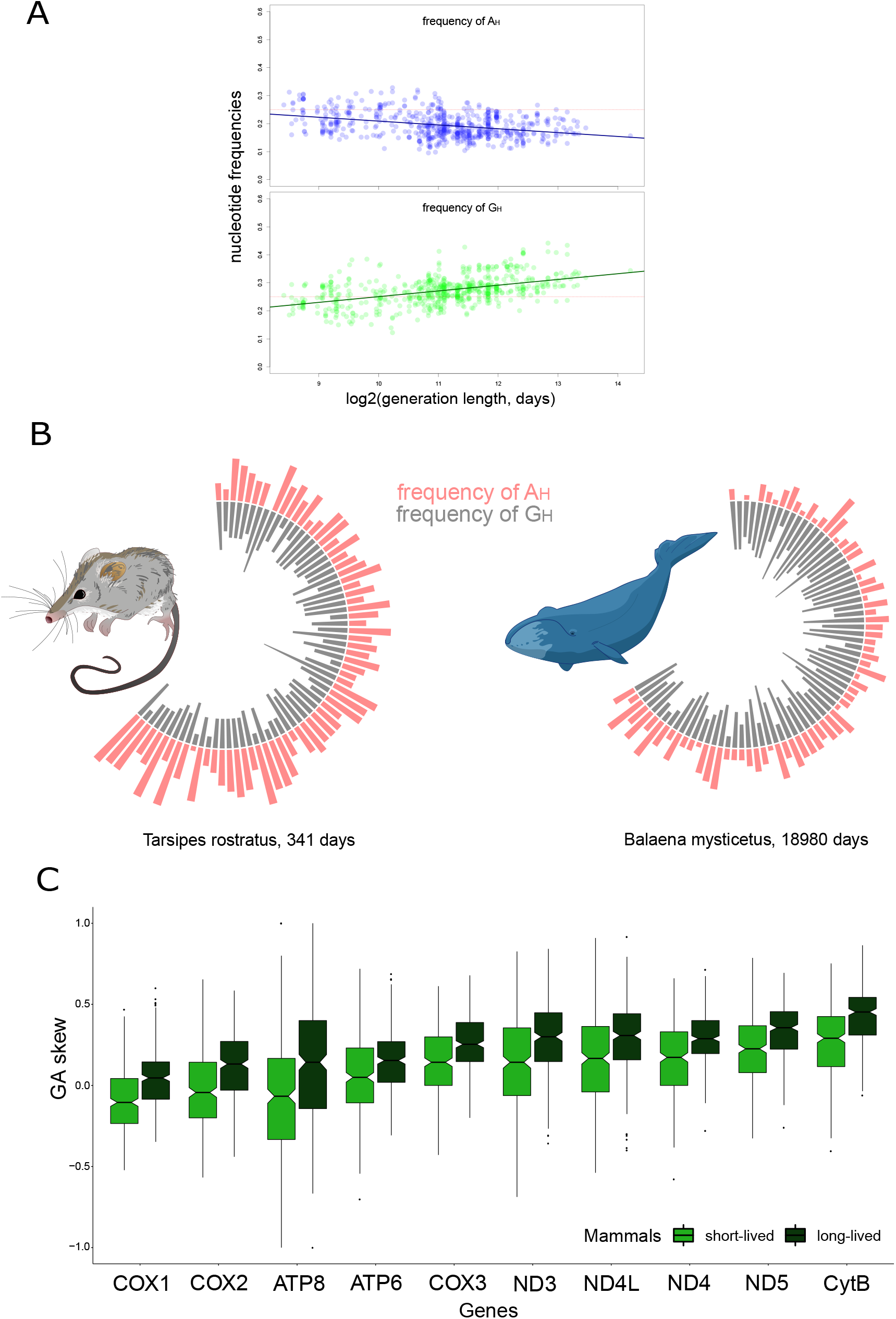
the long-term effect of the mutational bias: neutral nucleotide content in mammalian species. 3A. Nucleotide frequencies in neutral sites of all 13 protein-coding genes as a function of generation length - fraction of A_H_ is decreasing while fraction of G_H_ is increasing (N = 650). 3B. Frequencies of A_H_ (red) and G_H_ (grey) nucleotides along the major arc of mtDNA of the most short-lived and most long-lived species from our dataset. Each bar represents the nucleotide frequency in a 20 nucleotide window. In both mammals A_H_ is decreasing and G_H_ is increasing along the mtDNA: from bottom left (origin of replication of light strand) to top right (origin of replication of heavy strand). Also, mtDNA of a long-lived mammal has a deficit of A_H_ and excess of G_H_ genome-wide - a signature of increased longevity. 3C. Changes in nucleotide content along mtDNA of short- and long-lived mammals. (N = 650). All genes (except for ND6) located in the major arc are ranked according to the time spent single stranded: from COX1 to CYTB. Pairs of boxplots for each gene represent G_H_A_H_ skew for short- and long-lived mammals splitted by the median generation length.

An excess of G_H_ and deficit of A_H_ in long-lived species determines the G_H_A_H_ nucleotide skew which approximates the level of asymmetry in the distribution of these two nucleotides on heavy chain of mtDNA and is calculated as (G_H_-A_H_)/(G_H_+A_H_). As expected we observed positive correlation between G_H_A_H_ skew and generation length of mammalian species (phylogenetic generalized least squares: coefficient = 0.13, p-value = 2.9*10^−4^; see also Figure 3C). To visualize a contrast in G_H_A_H_ skew between the shortest- and the longest-lived species we plotted A_H_ and G_H_ fractions along the major arc of mtDNA for opossum (generation length 341 days) and whale (generation length 18980 days) (Figure 3B). An excess of G_H_ and deficit of A_H_ were more pronounced in whale versus opossum.

In parallel with the effect of generation length (Figure 3A), the frequencies of G_H_ and A_H_ are changing along the major arc (Figure 3B), which is reflected in the change in G_H_A_H_ skew with gene position. It has been shown that the frequency of A_H_>G_H_ substitutions depends on Time which heavy strand Spent in Single-Stranded (TSSS) condition during asynchronous mtDNA replication (Faith and Pollock 2003). Genes, located close to the origin of light strand replication (O_L_), such as *COX1*, spend minimal time being single-stranded and demonstrate the low frequency of A_H_>G_H_, while genes located far away from O_L_ spend more time being single-stranded and demonstrate correspondingly higher frequencies of A_H_>G_H_. Thus, we expect that G_H_A_H_ skew is a function of both: species-specific generation length and gene-specific TSSS. Plotting G_H_A_H_ skew of short- and long-lived mammals for each gene we observed that indeed G_H_A_H_ skew increases with both species-specific generation length and gene-specific TSSS (Figure 3C).

Increase in G_H_A_H_ skew with TSSS is supported by corresponding decrease in A_H_ and an increase in G_H_ with TSSS in all mammalian species (Figure 3A, Supplemental Mat 4.2). Interestingly, the rate of changes in the nucleotide frequencies (decrease in A_H_ and increase in G_H_) with TSSS is higher in long- than short-lived mammals (Supplemental Mat 4.2, the same effect is visible on Figure 3B). Faster changes (stronger gradients) in A_H_ and G_H_ along the genome of long-lived mammals can be interpreted as an interaction between TSSS and longevity probably mediated by the significantly increased oxidative damage of mtDNA with high TSSS in long-lived mammals.

Altogether, our results demonstrate that the neutral nucleotide content reflects the mutational bias from A_H_ to G_H_: long-lived mammals are more A_H_ poor and G_H_ rich (Figure 3A), have stronger G_H_A_H_ skew (Figure 3B, 3C) and faster changes in nucleotide content (decrease in A_H_ and increase in G_H_) along the genome (Supplemental Mat 4.2).

## Discussion

We observed that the fraction of A_H_>G_H_ substitutions increases with cellular and organismal longevity on different time scales - from several years in somatic tissues (chapter 1, Figure 1) and dozens of years in the human de novo germline substitutions (chapter 2) to hundreds of thousands years (the average time of segregation of neutral mtDNA polymorphisms (Atkinson et al. 2008)) in within-species polymorphisms (chapter 3, Figure 2) and millions of years as in the case of a comparative species scale (chapter 4, Figure 3).

We propose that a process of mtDNA mutagenesis, rather than selection, is primarily responsible for all our findings. Due to little or no evidence of selection (i) on mtDNA somatic mutations in human cancers (Yuan et al. 2020; Ju et al. 2014), (ii) on low-heteroplasmy de novo mutations in the germline (Arbeithuber et al. 2020) and (iii) on synonymous four-fold degenerate sites in mtDNA of mammals (Faith and Pollock 2003; Uddin and Chakraborty 2017) we consider the vast majority of variants analysed in this study as effectively neutral and thus driven predominantly by mutagenesis.

We propose that mtDNA mutagenesis is sensitive to the mitochondrial microenvironment which in turn depends on cell- and organism-specific longevity. We assume that oxidative damage - a main by-product of the aerobic metabolism - through the deamination of adenosine induces A_H_>G_H_ substitutions on a single-stranded mtDNA heavy strand during replication. Importantly, the well-documented ROS-induced 7,8-Dihydro-8-oxo-20-deoxyguanosine (8-oxodG), leading to G > T transversions, was expected to be one of the strongest mtDNA-markers of oxidative damage. However, mtDNA G > T is low frequency, age-independent and demonstrates weak if any association with oxidative damage in previous works (Yuan et al. 2020; Kennedy et al. 2013; Hoekstra et al. 2016; Zsurka et al. 2018) and our own results (chapters 1, 3 and 4). Of note G_H_>T_H_ and complementary to it C_H_>A_H_ are similar to each other, probably reflecting low mutagenic asymmetry, and demonstrate opposite to A_H_>G_H_ trend in our analyses (Figure 2C, Supplemental Mat 3.4) probably reflecting a weak effect of ROS on double stranded mtDNA. Here, instead of G>T, weakly affecting double-stranded DNA, we describe a new and strong mtDNA-marker of oxidative damage affecting single stranded DNA: A_H_>G_H_ substitution. Numerous recent papers, discussed below, support that A>G is a hallmark of oxidative damage of single-stranded DNA.

First, it has been shown for example that colon cancer, in the course of transformation to a more hypoxic stage, decreases its mtDNA mutation rate due to slowing down both common transitions: C_H_>T_H_ and A_H_>G_H_ (Ericson et al. 2012). In our study, we partially confirmed findings of Ericson extending it to all cancer types and claiming that hypoxic conditions, more often observed in fast-dividing cells and late cancer stages, are associated with decreased fraction of A_H_>G_H_ transitions.

Second, recent deep sequencing of de novo mtDNA mutations in aged versus young mice oocytes confirmed that an increased fraction of A_H_>G_H_ substitutions is the best hallmark of oocyte aging (Arbeithuber et al. 2020), which is in line with our trend observed in human mother-offspring duos (chapter 2). This discovery also suggests that continuous mtDNA turnover within the dormant oocytes can be a universal trait of all mammalian species (Arbeithuber et al. 2020) and thus variation in mtDNA mutational spectra between different species can reflect the species specific aging, dependent on oxidative damage in oocytes. Extending the results of Arbeithuber et al (Arbeithuber et al. 2020) from young and old mice oocytes to oocytes of short- and long-lived mammalian species we observed the same trend with A_H_>G_H_ marking more long-lived oocytes (Figure 2). These results suggest increased oxidative damage in mtDNA of oocytes of long-lived mammals.

Third, extension of the trend of the species-specific mtDNA mutational spectra (Figure 2) to nucleotide content of mitochondrial genomes (Figure 3) confirms long-term effects of the mutational spectrum and goes in line with described before excess of G_H_ in mtDNA of long-lived mammals (Lehmann et al. 2006, Lehmann et al. 2008). Although Lehmann et al proposed a selective explanation of this observation assuming increased stability of the G_H_-rich genomes which may confer an advantage to long-lived mammals, our findings demonstrate that an excess of G_H_ in long-lived mammals may be a result of mutagenesis, not selection.

Fourth, it is important to mention that A_H_>G_H_ transitions are the substitution type which is the most sensitive to the time being single-stranded during asynchronous replication of mammalian mtDNA (Faith and Pollock 2003; Tanaka and Ozawa 1994), which may be driven by the increased oxidative damage of single-stranded DNA.

Fifth, an excess of A:T>G:C has been observed in aerobically versus anaerobically grown Escherichia coli and the signal was driven by the lagging strand, spending more time in single-stranded condition (Shewaramani et al. 2017). Interestingly, A>G substitutions, associated with oxidative damage, can be a key process, explaining a long-standing evolutionary puzzle of increased GC content of aerobic versus anaerobic bacteria (Naya et al. 2002; Romero et al. 2009; Aslam et al. 2019).

Sixth, additional lines of evidence are coming from a mismatch repair pathway, which can prevent mutations caused by oxidative stress (Bridge et al. 2014). Mismatch repair-deficient mouse cell line shows a 6-50 fold increase in the rate of the A:T>G:C substitutions, and this increase is washed out in the presence of antioxidants (Shin and Turker 2002).

Seventh, it has been shown recently that the fraction of A_H_>G_H_ in mtDNA positively correlates with the ambient temperature of Actinopterygii species and it is higher in non-hibernating versus hibernating mammals (Mikhailova et al. 2020). Because higher temperature is associated with an increased level of aerobic metabolism (Martin and Palumbi 1993) we assume that A_H_>G_H_ can be a marker of oxidative damage.

Finally, A>G is the most asymmetric mutation in the nuclear genome, which is a hallmark of a mutagen, inducing strand-specific DNA damage (Seplyarskiy et al. 2019), which despite the fact that yet unknown, can be well mediated by oxidative damage.

The results of our study can significantly expand the usability of mtDNA mutational spectra. Low-heteroplasmy somatic mtDNA mutations from a merely neutral marker used to trace cellular lineages (Ludwig et al. 2019) can be transformed to a metric, associated with cell-specific aging state - a metric which can be especially important in highly heterogeneous tissues such as cancers. Low-heteroplasmy de novo germ-line mtDNA mutations (Arbeithuber et al. 2020) can predict the biological age of human oocytes, which can be used in *in vitro* fertilization techniques. MtDNA mutational spectra, reconstructed for non-model vertebrate species, may help to approximate their average generation length - a metric which is not always easy to estimate empirically.

Altogether, we propose that A>G substitutions is a new marker of the oxidative damage on the single-stranded DNA. Direct experimental investigation of the effect of different mutagens (Kucab et al. 2019) and especially an effect of oxidative damage on mutations (Degtyareva et al. 2019) and chemical modifications of nucleotides (Koh et al. 2018, Hao et al. 2020) on single-stranded mtDNA will significantly improve our understanding of mtDNA mutagenesis.

## Methods

### Analyses of somatic mtDNA mutations and mother-offspring pairs

We used 7611 somatic single-nucleotide mtDNA substitutions observed in human cancer samples (Yuan et al. 2017). Taking into account presumably very weak selection on somatic mtDNA mutations in cancers and their low VAFs we used all substitutions (not only synonymous fourfold degenerate) as neutral to derive the mutational spectrum. Correspondingly, to normalize the relative probabilities of mutations we used all nucleotides of the human genome (A, T, G, C) not just the ones in synonymous fourfold degenerate positions. Tissue-specific number of divisions of each stem cell per lifetime we sampled from the supplementary table 1 of (Tomasetti and Vogelstein 2015) and other sources (Supplemental Mat).

Data for the analyses of mother-offspring pairs were taken from three studies (Rebolledo-Jaramillo et al. 2014; Wei et al. 2019; Zaidi et al. 2019). In each dataset we filtered out all inherited mutations and took into account only de novo mtDNA mutations in offspring.

## Reconstruction of the species-specific mutational spectrum for mammalian species

Using all available intraspecies sequences (at April 2016) of mitochondrial protein-coding genes we derived the mutational spectrum for each species. Briefly, we collected all available mtDNA sequences of any protein-coding genes for any chordate species, reconstructed the intraspecies phylogeny using an outgroup sequence (closest species for analyzed one), reconstructed ancestral states spectra in all positions at all inner tree nodes and finally got the list of single-nucleotide substitutions for each gene of each species. The pipeline is described in more details in Supplementary Materials 3. Using species with at least 15 single-nucleotide synonymous mutations at four-fold degenerate sites we estimated mutational spectrum as a probability of each nucleotide to mutate into any other nucleotide (vector of 12 types of substitutions with sum equals one) for more than a thousand of chordata species. We normalized observed frequencies by nucleotide content in the third position of four-fold degenerative synonymous sites of a given gene. To eliminate the effect of nonuniform sampling of analyzed genes between different species the vast majority of our analyses were performed with *MT-CYB* gene - the most common gene in our dataset.

Generation length in days (as the average age of parents of the current cohort, reflecting the turnover rate of breeding individuals in a population) for mammals was downloaded from Pacifici et al. (Pacifici et al. 2013).

## Analyses of complete mitochondrial genomes

We downloaded whole mitochondrial genomes from GenBank using the following search query: “Chordata”[Organism] AND (complete genome[All Fields] AND mitochondrion [All Fields] AND mitochondrion[filter]. We extracted non-overlapping regions of protein-coding genes, calculated codon usage and extracted fractions of A_H_, T_H_, G_H_ and C_H_ nucleotides in synonymous fourfold degenerate positions. For each protein-coding gene of each reference mitochondrial genome of mammalian species we estimated G_H_A_H_ skew as (G_H_-A_H_)/(G_H_+A_H_) using only synonymous fourfold degenerate sites.

All statistical analyses were performed in R.

## Acknowledgments

We thank Han Liang, Peter Campbell and Yuan Yuan for the sharing of somatic mtDNA mutations from cancer samples, Athanasios Kousathanas for statistical comments, the whole laboratory of Alexandre Reymond, Mikhail Gelfand and Vladimir Katanaev for discussions and valuable suggestions. K.P. was supported by the 5 Top 100 Russian Academic Excellence Project at the Immanuel Kant Baltic Federal University. E.O.T. is supported by a scholarship from the Austrian Science Fund (FWF, DOC 33-B27). This work was also supported by Russian Foundation of Basic Research [No. 18-29-13055 &, 18-04-01143 to K.P; No. 19-29-04101 to I.M].

## Notes

### Competing Interest Statement

The authors have declared no competing interest.

### Summary of Updates

the paper is rewritten in a more clear style and all technical details are in supplements now

