## Supplementary materials for "Mammalian mitochondrial mutational spectrum as a hallmark of cellular and organismal aging"

Supplementary Materials to the manuscript by Mikhailova et al. “Mammalian mitochondrial mutational spectrum as a hallmark of cellular and organismal aging”

### (1) mtDNA mutational spectrum in human cancers changes during tumorigenesis and is associated with cell turnover rate in ancestral tissues

(1.1) mtDNA mutational spectrum is changing during tumorigenesis

https://github.com/polarsong/mtDNA_mutspectrum/blob/Cancer/Head/2Scripts/Cancer.TimeOfTumorigenesis.R

MtDNA somatic mutations observed in human cancer cells can represent two populations: (i) early mutations which occurred before the tumorigenesis and expanded later by clonal expansion of cancer cells and (ii) late mutations, originated in cancer cells. To split all somatic mutations into presumably early and presumably late we used variant allele frequencies (VAF). First, we compared VAF in matched cancer and normal tissues and observed positive correlation between them (Spearman's rho = 0.07, p = 1.226e-09 for the whole subset of substitutions including non-observed, i.e. zero VAF, in normal tissue N = 7611; Spearman's rho = 0.09, p = 9.638e-11 for a subset of variants with non-zero VAF in normal tissues, N = 5436) suggesting that many mtDNA somatic mutations indeed originated before the tumorigenesis. Second, we compared cancer specific VAFs between samples with (N=5436) and without (N=2265) corresponding mutations in normal tissues. As expected, we observed increased cancer-specific VAFs in samples, where the same mutation, even with low VAF has been observed in normal tissue (Figure S1A, p = 0.005, Mann-Whitney U-test). Altogether these analyses confirm that we can approximate the time of origin of mutations using VAF: high VAF marks mutations originated early and low VAF marks late mutations.

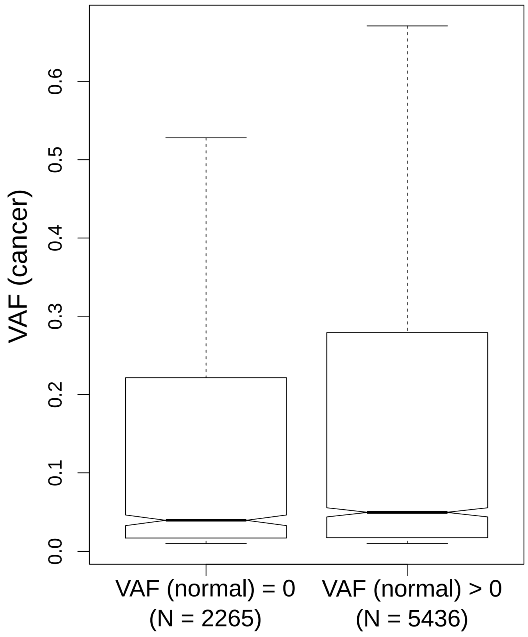


*Figure S1A. Variant Allele Frequency (VAF) in cancer approximates the time of origin of the variant: it is high for early variants (observed also in normal tissues, N = 5436) and it is low for late variants (not observed in normal tissues, N = 2265).*

Next using cancer VAF as an approximation for the time of origin of the variant we wanted to observe how mtDNA mutational spectra is changing during tumorigenesis.

First, we have split all variants by the median value of VAF (4.5%) and estimated Ts/Tv for variants with low and high VAFs as 10.3 and 14.9 correspondingly (Fisher Odds Ratio = 0.69, p = 2.524e-05, N = 7611, figure S1B left panel). This result is robust to an elimination of potentially low-quality variants: when we removed 25% of variants with the lowest p-values (variants with -log10(p-value) <= 59) we saw nearly identical result (median of VAF = 9.4%, Fisher Odds Ratio = 0.67, p = 0.0002, N = 5709). Both common transitions $A_H>G_H$ and $C_H>T_H$ contribute independently and significantly to this trend: $A_H>G_H$/Tv: Fisher Odds Ratio = 0.68, p = 5.26e-05; $C_H>T_H$/Tv: Fisher Odds Ratio = 0.72, p = 0.0002.

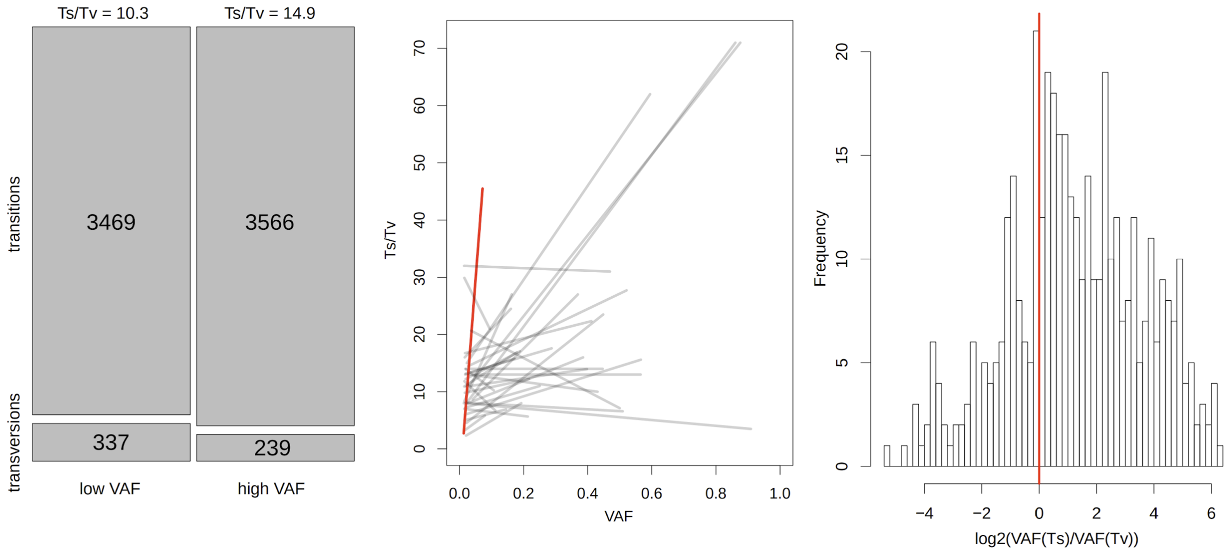


*Figure S1B. Ts/Tv is higher for early (with high VAF) versus late (with low VAF) variants. The integral analysis includes all somatic variants (left panel). Cancer-type specific analysis (middle panel) takes into account the trend specific for each cancer type; each cancer type is marked by a segment, PBCA is marked by the red segment. The sample-specific analysis demonstrates increased VAF(Ts) versus VAF(Tv) within each sample (right panel).*

Second, to rule out the potential effect of different cancer types on our integral analysis (Figure S1B left panel), we tested if there is an increase in Ts/Tv with VAF for each cancer type individually (Figure S1B middle panel). To do it within each cancer type we split all variants into rare and common (by the median VAF) and calculated for each group Ts/Tv and mean VAF. We expected that with the increase of VAF, Ts/Tv would also grow. To visualize it we plot segments for each cancer type connecting Ts/Tv and VAF or rare variants with Ts/Tv and VAF of common variants. Altogether these segments demonstrate positive trend (p = 0.0006, paired Mann-Whitney U-test, N = 36 cancer types, figure S1B middle panel), however, individually only Pediatric Brain Cancer (PBCA) demonstrated a significant increase in Ts/Tv with VAF (median of VAF = 10.1%, Fisher Odds Ratio = 0.06, p = 1.14e-06, N = 186, it is marked by red line on figure S1B, middle panel). Elimination of PBCA from the dataset doesn't affect results (p = 0.001, paired Mann-Whitney U-test, N = 35 cancer types). The same trend is also supported by both common transitions independently: $A_H>G_H$/Tv (p = 0.0004, paired Mann-Whitney U-test, N = 36) and $C_H>T_H$/Tv (p = 0.0006, paired Mann-Whitney U-test, N = 36).

Third, to rule out additional confounders such as different purity of samples, we tested if there is an increase in Ts/Tv with VAF within each sample. To do this we used a subset of samples with at least one transition and at least one transversion (N = 419) and estimated mean VAF for Ts and Tv in each sample. Comparing them within each sample we observed higher VAF for Ts than for Tv (p = 5.109e-15, Mann-Whitney Paired test, N = 419) and correspondingly log ratios of VAF(Ts)/VAF(Tv) was higher than expected value of zero (median of the log ratios of Ts/Tv is 1.1665; p-value < 2.2e-16, Wilcoxon test with mu = 0; figure S1B right panel). Both common transitions contribute significantly to this trend: $A_H$>$G_H$/Tv ($p$ =1.501e-06, Mann-Whitney Paired test, N = 286) and $C_H$>$T_H$/Tv ($p$ = 2.651e-09, Mann-Whitney Paired test, N = 346).

In order to analyze the contribution of $A_H$>$G_H$ versus $C_H$>$T_H$ we compared VAF($A_H$>$G_H$) and VAF($C_H$>$T_H$) within each sample and found out that $A_H$>$G_H$ is characterized by slightly higher VAF as compared to $C_H$>$T_H$ (median of the differences between VAF ($A_H$>$G_H$) and VAF($C_H$>$T_H$) is 0.1%, $p$ =0.0097, Mann-Whitney paired test, N = 983), meaning that $A_H$>$G_H$ on average happens more early on than $C_H$>$T_H$. Because the relationship between these two most common transitions doesn't depend on rare transversions, this result independently confirms robustness of our conclusion about the changes in the pattern of mutagenesis during tumorigenesis.

Putting together all these analyses: the integral one (figure S1B, left panel), the cancer-type-specific one (figure S1B, middle panel) and sample-specific one (figure S1B, right panel), we conclude that mutational spectrum is changing during tumorigenesis - becoming less transition rich and especially less $A_H$>$G_H$ rich in more advanced cancers.

(1.2) mtDNA mutational spectrum differs between cancer types

https://github.com/polarsong/mtDNA_mutspectrum/blob/Cancer/Head/2Scripts/Cancer.DifferencesBetweenCancerTypes.R

Using an approximate turnover rate of stem cell divisions in each of 21 normal tissue samples, ancestral to our analyzed cancer types we split all cancer tissues into three categories: fast-replicating - with stem cell turnover rate less than a month (uterus, colon/rectum, stomach, cervix, esophagus, head/neck, lymphoid and myeloid tissues), intermediate - with stem cell turnover rate higher than a month and less than ten years (breast, prostate, skin, bladder, biliary, pancreas, liver, and kidney) and slow-replicating - with stem cell turnover rate higher than ten years (thyroid, lung, bone/soft tissue, CNS, ovary). The table S1 provides estimations of stem cells turnover rates, used in downstream analyses.

*Table S1. Turnover of stem cells in different human tissues  (in days)*

| Cancer Tissue : GTEx tissue annotation | Turnover Rate of stem cells (in days) | Source |
|---|---|---|
| Bladder : Bladder | 200.0 | (Wang, Ross, and Mysorekar 2017), (“Urothelial Stem Cell Regeneration” n.d.). |
| | 49 | (Seim, Ma, and Gladyshev 2016) |
| Bone/SoftTissue | 5373.0 | (Tomasetti and Vogelstein 2015) |
| Breast | 84.5 | (Tomasetti et al. 2017) |
| Biliary | 400.0 | Here we use the same turnover rate as in liver (Tomasetti and Vogelstein 2015) |
| Cervix | 6.0 | (“Bionumbers” n.d.) |
| Lymphoid | 30.0 | (Tomasetti and Vogelstein 2015) |
| Myeloid | 30.0 | (Tomasetti and Vogelstein 2015) |
| Colon/Rectum | 5.0 | (Tomasetti and Vogelstein 2015) |
| | 3.5 | (Seim, Ma, and Gladyshev 2016) |
| Prostate | 120.0 | (Tomasetti et al. 2017) |
| Esophagus | 11.0 | (Tomasetti et al. 2017) |
| | 10 | (Seim, Ma, and Gladyshev 2016) |
| Stomach | 5.5 | (“Bionumbers” n.d.) |
| CNS | 10000.0 | (Tomasetti and Vogelstein 2015) |
| | 32850 | (Seim, Ma, and Gladyshev 2016) |
| Head/Neck | 16.0 | (Tomasetti and Vogelstein 2015) |
| Kidney | 1000.0 | (Yue Li 2013) |

| | | |
|---|---|---|
| | 270 | (Seim, Ma, and Gladyshev 2016) |
| Liver | 400 | (Tomasetti and Vogelstein 2015) |
| | 327 | (Seim, Ma, and Gladyshev 2016) |
| Lung | 5143 | (Tomasetti and Vogelstein 2015) |
| | 200 | (Seim, Ma, and Gladyshev 2016) |
| Ovary | 11000 | (Tomasetti and Vogelstein 2015) |
| Pancreas | 360 | (Tomasetti and Vogelstein 2015) |
| Skin | 147 | (Tomasetti and Vogelstein 2015) |
| | 64 | (Seim, Ma, and Gladyshev 2016) |
| Thyroid gland | 4138.0 | (Tomasetti and Vogelstein 2015) |
| | 3180 | (Seim, Ma, and Gladyshev 2016) |
| Uterus | 4 | (Kim, Tavaré, and Shibata 2005) |
| Heart muscle | 25300 | (Seim, Ma, and Gladyshev 2016) |
| Skeletal muscle | 5510 | (Seim, Ma, and Gladyshev 2016) |
| Male gametes | 60 | (“Bionumbers” n.d.) |
| Small intestine epithelium | 2-4 | (“Bionumbers” n.d.) |

Using all somatic mutations observed in the cancer samples belonging to three groups we estimated their Ts/Tv as 10.0, 12.6 and 14.5 correspondingly for fast, intermediate and slow-dividing tissues. To test if the observed difference is robust, we estimated Ts/Tv for each groups after a thousand 50% jackknife re-samplings of mutations, and confirmed a trend of the increased Ts/Tv in slow- versus fast-dividing groups of tissues (Figure S1C left panel; all boxplots are generated on the basis of thousand 50% jackknife re-samplings). Both common transitions $C_H>T_H$ and $A_H>G_H$ independently contribute to this trend ($A_H>G_H$/Tv: 2.8, 3.8, 4.6; $C_H>T_H$/Tv: 5.0, 6.6, 7.5 for fast, intermediate and slow-replicating tissues, Figure S1C left panel).

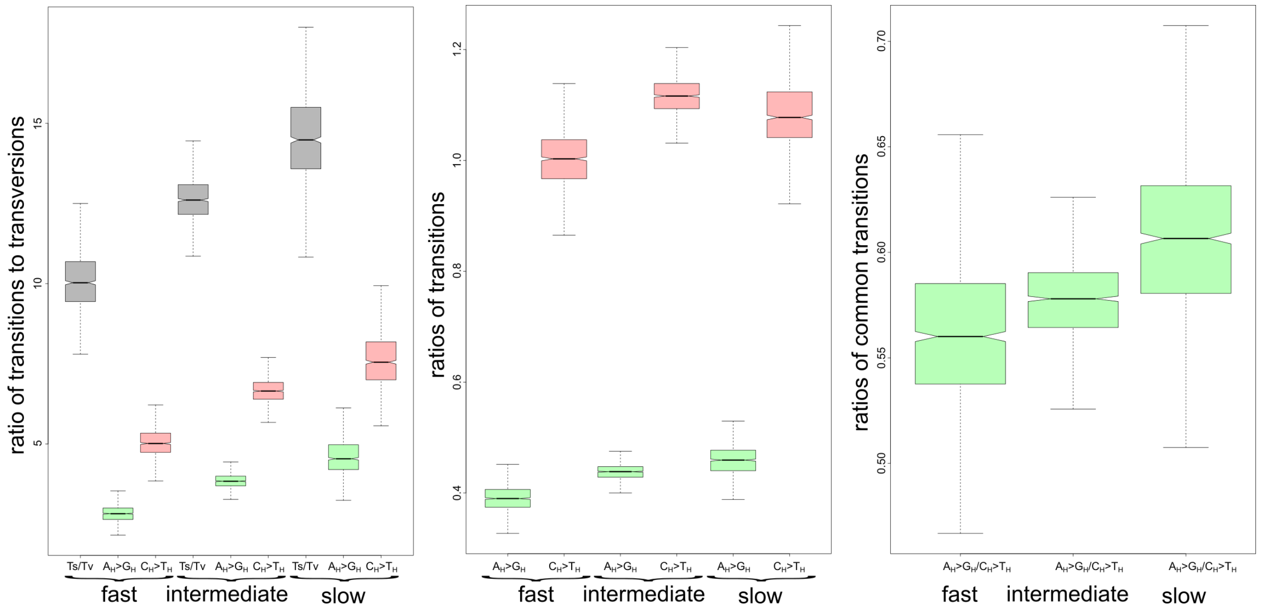


*Figure S1C. Slow-dividing tissues are enriched in transitions, primarily in $A_H$>$G_H$. Left panel: all substitutions normalized by Tv - Ts/Tv, $A_H$>$G_H$/Tv and $C_H$>$T_H$/Tv are higher for cancer types, descendant from slow versus fast-replicating tissues. Middle panel: all substitutions normalized by transitions - only $A_H$>$G_H$, but not $C_H$>$T_H$ demonstrates the concordant increase in slow-replicating tissues. Right panel: all substitutions normalized by the most common transition $C_H$>$T_H$ - $A_H$>$G_H$/$C_H$>$T_H$ is increasing from fast to slow-replicating tissues. All nucleotides on the plot are in heavy strand notation. The plotted distributions are derived based on thousand 50% jackknife re-samplings of mutations from corresponding cancer samples.*

Since transversions are very low frequency variants and some of them might be sequencing errors (Chen et al. 2017) we wanted to see whether this trend would still be visible when analyzing only transitions. We estimated the fraction of $A_H$>$G_H$ to all other transitions ($C_H$>$T_H$, $T_H$>$C_H$, $G_H$>$A_H$) as well as fraction of $C_H$>$T_H$ to all other transitions ($T_H$>$C_H$, $A_H$>$G_H$, $G_H$>$A_H$) and observed a corresponding increase in the fraction of $A_H$>$G_H$ ($A_H$>$G_H$/Ts: 0.39, 0.44, 0.46) but not in the fraction of $C_H$>$T_H$ ($C_H$>$T_H$/Ts: 1.00,1.12, 1.08) (Figure S1C middle panel). Finally, we compared the effects of these common transitions with each other and observed an increase in $A_H$>$G_H$ fraction over $C_H$>$T_H$ in slow- versus fast-dividing tissues ($A_H$>$G_H$/$C_H$>$T_H$: 0.56, 0.58, 0.61 for fast-, intermediate and slow-replicating tissues, Figure S1C right panel). We conclude that the fraction of $A_H$>$G_H$ is increasing from fast- to slow-replicating tissues.

(1.3) mtDNA mutational spectrum as a function of both VAF and tissue-specific turnover rate

*Table S2. Results of multiple logistic regressions linking the binomial variable* $A_H>G_H$ *(1 = yes or 0 = no) with independent variables: tissue-specific cell turnover rate; variant allele frequency (VAF) of each mutation in cancer sample; mtDNA copies; patient age. The results were obtained with glm function in R with "family = binomial" setting. All variables were scaled.*

| Model | Coefficient | Variable | p-value |
|---|---|---|---|
| 1A) main model, visualized in the figure 1 of the main text. Dependent variable is 1 ($A_H>G_H$) or 0 (any other substitutions), N = 7611. | -0.95 | intercept | < 2e-16 |
| | 0.049 | cell turnover in days | 0.049 |
| | 0.088 | VAF | 0.0004 |
| 1B) identical to the model 1A, but continuous variable "cell turnover in days" is substituted by Dummy variable which equals 1 in case of fastly dividing samples and 0 in case of intermediate and slow dividing samples. | -0.95 | intercept | < 2e-16 |
| | -0.065 | Dummy (fastly-dividing) | 0.013 |
| | 0.093 | VAF | 0.0002 |
| 1C) identical to the model 1B, but analyzed mutations are moderately common (VAF > 0.01738204, which is the lower quartile of the VAF distribution), N = 5708. | -0.94 | intercept | < 2e-16 |
| | *-0.079* | Dummy (fastly-dividing) | *0.0089* |
| | *0.102* | VAF | *0.0004* |
| *1D) identical to the model 1B, but analyzed mutations are transitions only. N = 7035.* | *-0.842* | intercept | < 2e-16 |
| | *-0.054* | Dummy (fastly-dividing) | *0.043* |
| | *0.071* | VAF | *0.0058* |
| *2A) identical to the model 1A, but with two additional variables: mtDNA copies and patient age, N = 7611.* | *-0.95* | intercept | < 2e-16 |
| | *0.057* | cell turnover in days | *0.029* |
| | *0.087* | VAF | *0.0006* |
| | *-0.024* | *patient age* | *0.367* |
| | *-0.076* | *mtDNA copies* | *0.004* |
| *2B) identical to the model 2A, but with removed non-significant variable 'patient age', N = 7611.* | *-0.955* | intercept | *< 2e-16* |
| | *0.0578* | cell turnover in days | *0.0217* |
| | *0.090* | VAF | *0.0003* |
| | *-0.068* | *mtDNA copies* | *0.0090* |

(1.4) mtDNA mutational spectrum in cancer and hypoxia

https://github.com/polarsong/mtDNA_mutspectrum/blob/Cancer/Head/2Scripts/Cancer.Hypoxia%26MutSpec.R

**(2) mtDNa mutational spectrum is changing with female reproductive age**

Analysis of data from Rebolledo et al (Rebolledo-Jaramillo et al. 2014; Wei et al. 2019; Zaidi et al. 2019). $A_H$>$G_H$ is characterized by the highest age of fertilization (average age of fertilization is 34.1, N = 4) as compared to $C_H$>$T_H$ (average age of fertilization is 29.9, N = 4) or all other substitutions (average age of fertilization is = 32.1, N = 12). Due to the small sample size (16 mutations in total) this trend was not significant (p = 0.19 if we compare $A_H$>$G_H$ with all other substitutions and p = 0.0956 if we compare $A_H$>$G_H$ with $C_H$>$T_H$; one-sided Mann-Whitney U test).

GitHub:
https://github.com/polarsong/mtDNA_mutspectrum/blob/Humans/Head/2Scripts/Humans.RebolledoAnalyses.R

Analysis of data from Wei et al (Wei et al. 2019). Data on the age of reproduction were not available and we used VAF of each de novo mtDNA variant as a function of oocyte age. We expect that $A_H$>$G_H$ mutations occur mainly in aged oocytes and thus are characterized by lower VAF as compared to other substitutions. To decrease potential selection effects we concentrated only on rare (with VAF < 10%) substitutions. Next, subsetting offspring with at least two rare de novo mtDNA transitions, one being $A_H$>$G_H$ and another non $A_H$>$G_H$, (23 offspring in total) we observed that $A_H$>$G_H$ on average has 1% lower VAF as compared to non $A_H$>$G_H$. The statistical difference between VAF($A_H$>$G_H$) and VAF(non $A_H$>$G_H$) was marginally significant (p = 0.05015, paired one-sided Mann-Whitney U test).

GitHub:
https://github.com/polarsong/mtDNA_mutspectrum/blob/Humans/Head/2Scripts/Wei2019OffspringMutTypesComparison.R

Analysis of data from Zaidi et al (Rebolledo-Jaramillo et al. 2014; Wei et al. 2019; Zaidi et al. 2019). Analyzing 31 children with at least one $A_H$>$G_H$ mutation and 66 children with at least one another transition, we observed that the median value of the age of fertilization is higher for children with at least one $A_H$>$G_H$ mutation (31.41 compared to 30.61 for other substitutions).

GitHub: https://github.com/polarsong/mtDNA_mutspectrum/blob/Humans/Head/2Scripts/Humans.ZaidiAnalyses.R

**(3) Mutational spectrum, derived from the intra-species mtDNA polymorphisms of mammalian species, is shaped by generation time**

(3.1) Pipeline to derive mtDNA mutational spectra for mammalian species

The most important pipeline stages were the following. After the retrieval of all available nucleotide sequences of mitochondrial protein-coding genes we generated local nucleotide BLAST-database. We retrive intraspecies sequences from this database using *tblastn* software (ncbi-blast 2.6.0+ package) and RefSeq protein query sequence of given species and given mitochondrial gene. At the next stage, all intraspecies sequences for given species and for given gene were aligned multiple times using *macse* v1.01b. This software performs codon-based multiple alignment (we used standard mitochondrial genetic code), and allows to check for inner stop codons in analyzed sequences (we select out such positions from alignment). Next, we reconstructed the ancestral sequences in each inner tree node. The reconstruction of intraspecies ancestral sequences was based on maximum likelihood tree topology and was made using two alternative approaches: maximum parsimony and maximum likelihood. Unrooted tree topology of intraspecies sequences was derived by *RaxML* v.8.2.9 and "-m GTRGAMMAIX" option, we used best tree from 50 alternative runs on distinct starting trees ("-N 50" option) for selecting best tree topology. After that tree was rooted by nearest neighbor sequence from other species. This nearest neighbor sequence was found as a first blast hit followed by sequences of given species in our sequence database. Maximum parsimony ancestral reconstruction was done using *dnapars* program of *phylip* v.3.697 package. Maximum likelihood reconstruction (marginal ancestral states reconstruction) was done using *RaxML* v.8.2.9 and "-m GTRGAMMAIX -f A" options. For ancestral reconstruction we used nucleotide-based (not codon based) substitution models due to only using only four-fold degenerate sites in codons for analyses in our work.

(3.2) Normalisation

The fractions of observed substitutions depend on the frequencies of ancestral nucleotides in the synonymous fourfold degenerate sites, which are highly non-uniform. After normalizing the observed nucleotide substitutions (Figure S2A left panel) by the ancestral nucleotide frequency (Figure S2A right panel) we obtained a mutational spectrum as a probability of a given nucleotide to mutate to any other nucleotide irrespective of its frequency. Hereafter in all downstream analyses of this chapter, we used these species-specific mutational spectra, presented as vectors of twelve probabilities with the total sum equal to one.

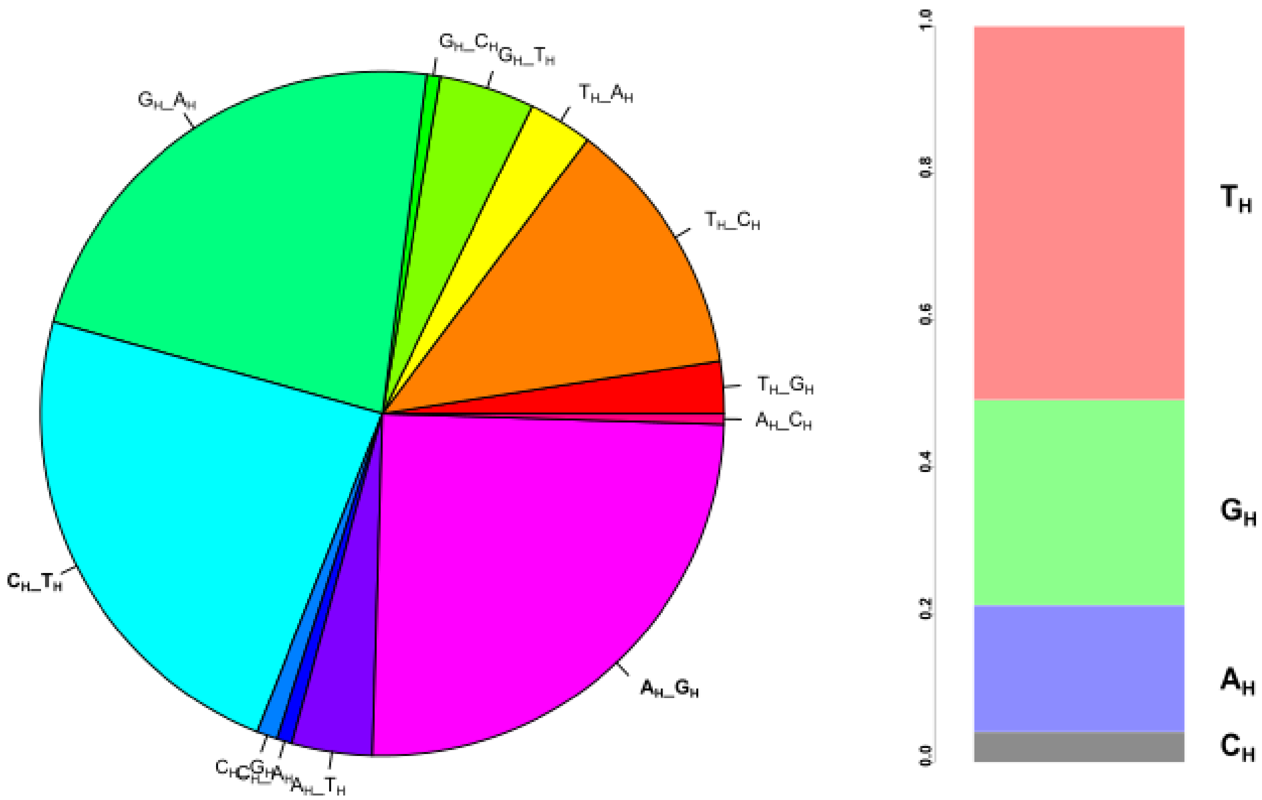


*Figure S2A. Derivation of mtDNA mutational spectrum for mammalian species (N = 611). Left panel: observed frequencies of twelve types of nucleotide substitutions in four fold degenerate synonymous sites of all available mtDNA protein-coding genes. Right panel: nucleotide content in four fold degenerate synonymous sites of all available mtDNA protein-coding genes.*

(3.3) species-specific Ts/Tv increases with generation length
(script: *VertebratePolymorphisms.MutSpecComparisons.Analyses.Ecology.Mammals.R*):

There is a positive correlation between Ts/Tv and species-specific Generation Length (Spearman's rho = 0.23, p-value = 2.021e-06, N = 424).

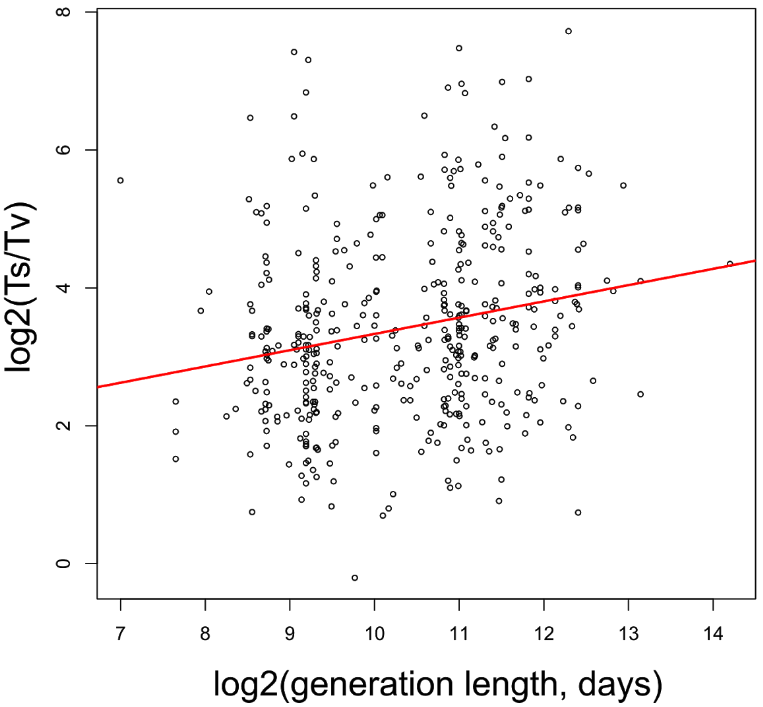


*Figure S2B. Positive correlation between generation length and Ts/Tv in mammals*

An increase in Ts/Tv together with generation length is being observed also when we split all mammals into several groups: (i) by quartiles of the generation length, (ii) by median of the generation length and (iii) by families.
When we split mammalian species into four quartiles, Ts/Tv is increasing from lower to higher quartiles: all possible pairwise comparisons of quartiles, except the comparison of the first and the second ones, show significant difference in Ts/Tv (Mann-Whitney U test, all p-values < 0.05).

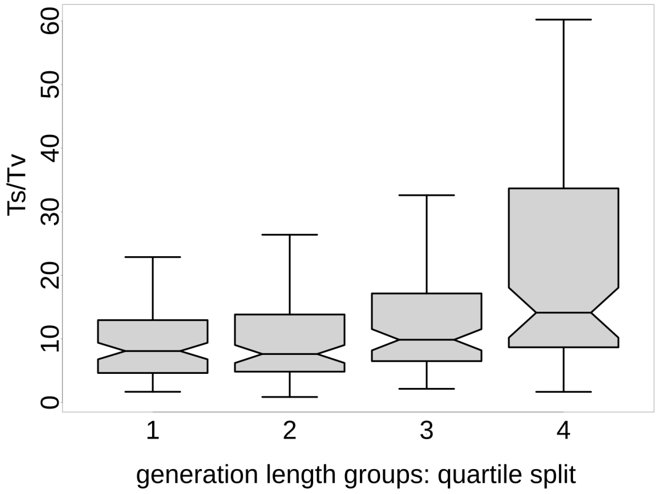


Median split of mammalian species by the generation length (median = 1497 days) also shows higher Ts/Tv in long- versus short-lived mammals (p-value = 1.687e-06, Mann-Whitney U test).

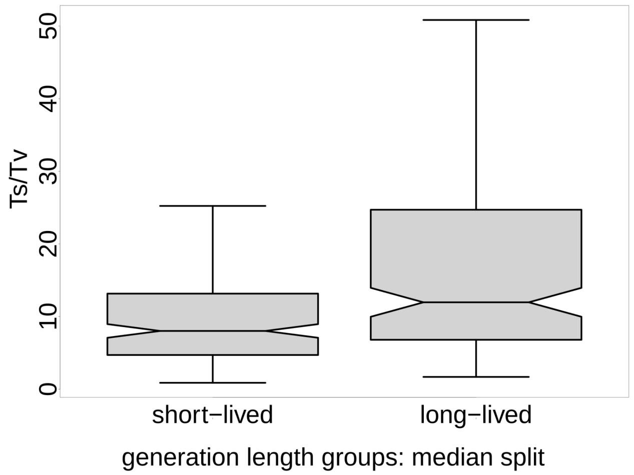


An increase in Ts/Tv with generation length is also pronounced on a family level. For each mammalian family, containing at least 3 species in our dataset, we estimated median Ts/Tv and median of generation length. Using Spearman rank correlation we confirmed a positive trend between these values ($p = 0.0016$, Spearman Rho = 0.85, N = 11). In the table below there are descriptive statistics of eleven families and double boxplots visualize distribution of Ts/Tv and generation length in each family.

| **Family** | **median Ts/Tv** | **median generation length (in days)** | **abbreviation** | **number of species** |
|---|---|---|---|---|
| Insectivora | 8.81 | 427 | Ins | 41 |
| Rodentia | 7.46 | 601 | Rod | 120 |
| Didelphimorphia | 7.08 | 643 | Did | 18 |
| Lagomorpha | 6.55 | 1047 | Lag | 19 |
| Chiroptera | 10.75 | 2064 | Chi | 81 |
| Carnivora | 13.21 | 2435 | Car | 34 |
| Cervidae | 13.33 | 2555 | Cer | 7 |
| Suidae | 10.57 | 2606 | Sui | 4 |
| Bovidae | 13.34 | 2870 | Bov | 16 |
| Primates | 14.14 | 3806 | Pri | 49 |
| Cetacea | 27.74 | 5158 | Cet | 6 |

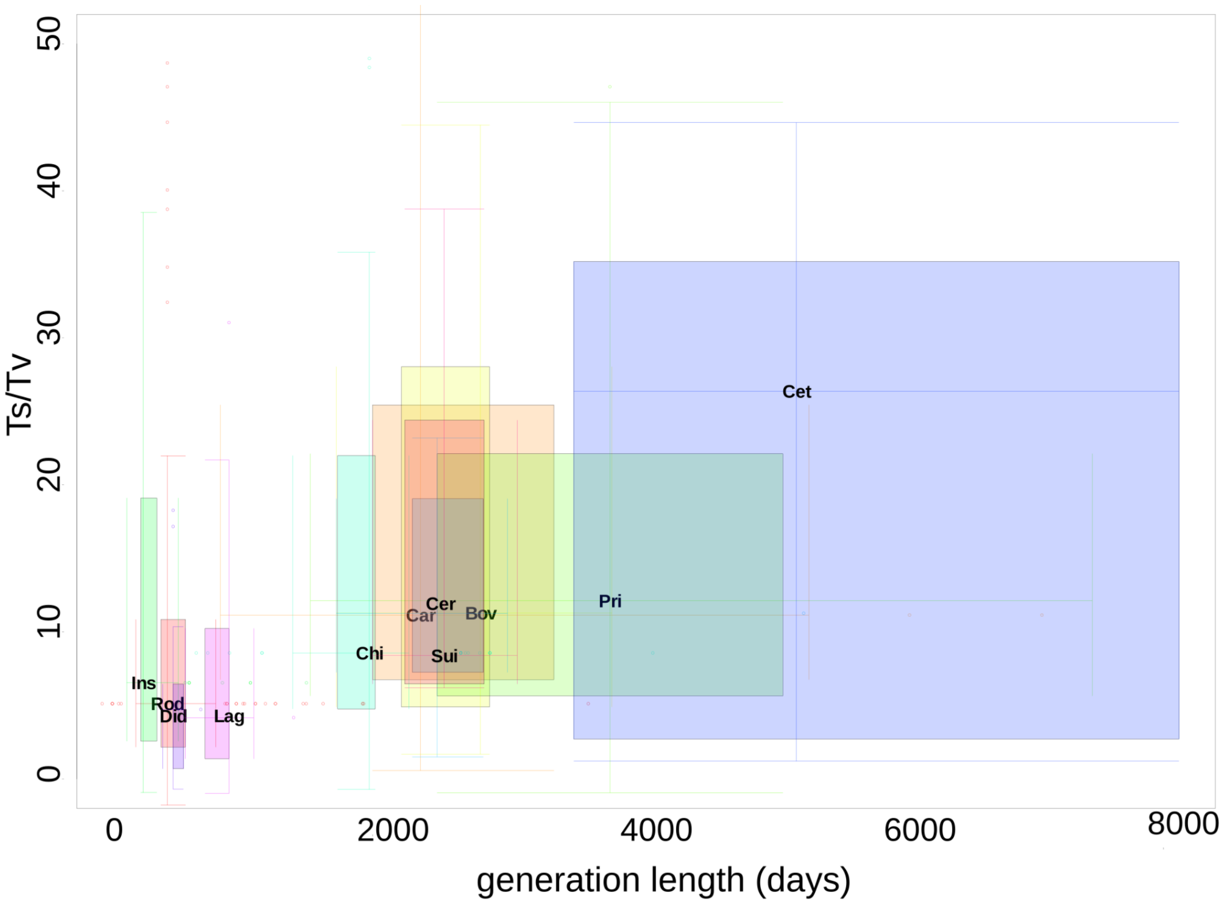


Boxplots were obtained using the R library "boxplotdbl". The filled surface for each family reflects interquartile range, cross of lines represents medians, whiskers mark minimum and maximum, excluding outliers (some outliers are not shown on the plot). A position of the three-letter abbreviation for each family corresponds to medians of the generation length and Ts/Tv of the corresponding family.

Although several analyses above demonstrated the robustness of the association between Ts/Tv and generation length we also performed phylogenetic generalised least-squares regression (PGLS) implemented in R library "caper". Due to the necessity to merge our dataset with an available phylogenetic tree we kept less than a half of our species (N = 211), but despite this fact the main trend is still significant. First, we demonstrated that Ts/Tv positively correlates with generation length (model 1A). Second, we added to the model the number of mutations as a number of reconstructed synonymous four-fold degenerate substitutions in CYB, which were used to reconstruct species-specific mutational spectrum (model 1B). Despite the fact that the number of mutations negatively correlates with Ts/Tv (probably due to higher sample size of short-lived mammals), the effect of the generation length on Ts/Tv stayed significant. Third, noting the non-significant deviation of intercept from the zero in model 1B we rerun the same model through the origin (fixing intercept at zero) and obtained strong and significant correlation between Ts/Tv and generation length (model 1C). In the final model 1C generation length is associated with Ts/Tv much more significant as compared to the 'number of mutations'.

| ***model*** | ***variable*** | ***coefficients*** | ***p values*** |
|---|---|---|---|
| ***1A: Ts/Tv ~ log2(generation length)****;* *N=211, lambda [ ML] = 0.000* | *intercept* | *-26.4047* | *0.094182* |
| | *log2(generation length)* | *4.1771* | *0.004472* |
| ***1B: Ts/Tv ~ log2(generation length)*** + | *intercept* | *12.2181* | *0.558976* |

| | | | |
|---|---|---|---|
| ***log2(number of mutations)**;* *N=211, lambda [ ML] = 0.000* | *log2(generation length)* | *2.9679* | *0.048715* |
| | *log2(number of mutations)* | *-4.3222* | *0.006407* |
| ***1C: Ts/Tv ~ 0 + log2(generation length) + log2(number of mutations)**;* *N=211, lambda [ ML] = 0.000* | *log2(generation length)* | *3.75742* | *2.454e-08* |
| | *log2(number of mutations)* | *-3.70511* | *0.001629* |

(3.4) fraction of $A_H$>$G_H$ increase in species with long generations (script: *VertebratePolymorphisms.MutSpecComparisons.Analyses.Ecology.Mammals.R*):

We included all five types of substitutions ($A_H$>$G_H$, $T_H$>$A_H$, $T_H$>$G_H$, $C_H$>$A_H$ and $G_H$>$T_H$ - see the main text, chapter 3) into the initial multiple linear model. Next, following the logic of the stepwise backward multiple model (removing one the most non-significant variable at each step) we determined that three types of substitutions were independently associated with generation length (*GL*).

*log2(GL) ~ 10.39 + 0.26*($A_H$>$G_H$) - 0.18*($T_H$>$A_H$) - 0.16*($C_H$>$A_H$),* equation (I)
*N = 424, $R^2$=0.104,*
*total p-value = 1.249e-10; p-values of transversions are < 0.002, p-value of $A_H$>$G_H$ = 6.42e-06,*
*presented coefficients are scaled.*

Comparing the scaled regression coefficients we observed that the strongest (the highest absolute effect size), as well as the most significant (the lowest p-value) effect was associated with $A_H$>$G_H$ transitions. Positive correlation of generation length with the frequency of a $A_H$>$G_H$ transition and negative correlation with the frequencies of $C_H$>$A_H$ and $T_H$>$A_H$ transversions as expected lead to the increased Ts/Tv in long-lived species described above.

It is important to note that the inclusion into the linear model (equation I) of the total number of mutations used to estimate the species-specific mutational spectrum (synonymous mutations in four fold degenerate sites in *MT-CYB* gene) does not affect these results quantitatively:

*log2(GL) ~ 10.39 + 0.24*($A_H$>$G_H$) - 0.14*($T_H$>$A_H$) - 0.17*($C_H$>$A_H$) - 0.26*(Number Of Mutations)* equation (Ia)
*N = 424, $R^2$=0.145,*
*total p-value = 2.464e-14; p-value ($A_H$>$G_H$) = 1.66e-05; p-value ($T_H$>$A_H$) = 0.0103; p-value ($C_H$>$A_H$) = 0.00249; p-value (Number Of Mutations) = 5.29e-06; presented coefficients are scaled.*
We can see that the number of mutations negatively correlates with GL, probably because short-lived, small-bodied mammals are better investigated and have more sequences deposited in GenBank. Importantly, all effects demonstrated in the chapter 3 of the main text and in the equation I stay qualitatively similar.

Similarly with PGLS analyses in suppl mat 3.3 we repeated three models, analyzing an association of $A_H$>$G_H$ with generation length. The final model 2C shows a significant positive effect of the generation length on $A_H$>$G_H$ and the absence of the effect of the 'number of mutations'

| *model* | *variable* | *coefficients* | *p values* |
|---|---|---|---|
| ***2A: $A_H$>$G_H$ ~ log2(generation length)**;* *N=211, lambda [ ML] = 0.336* | *intercept* | *0.0202212* | *0.8184* |
| | *log2(generation length)* | *0.0115248* | *0.1488* |

| | | | |
|---|---|---|---|
| ***2B: $A_H$>$G_H$ ~ 0 + log2(generation length)**; N=211, lambda [ ML] = 0.317* | *log2(generation length)* | *0.0133441* | *3.173e-08* |
| ***2C: $A_H$>$G_H$ ~ 0 + log2(generation length) + log2(number of mutations)**; N=211, lambda [ ML] = 0.329* | *log2(generation length)* | *0.0123511* | *0.0003097* |
| | *log2(number of mutations)* | *0.0018587* | *0.6942430* |

(3.5) PCA analyses: second PC represents a signature of a longevity associated mutagen (script: *VertebratePolymorphisms.MutSpecComparisons.Analyses.Ecology.Mammals.R*):

The correlation of the second principal component with generation length was significantly higher (rho = -0.39, p < 2.2e-16, N = 424) as compared to the sole effect of $A_H$>$G_H$ (Spearman's rho = 0.252, p value = 1.188e-07, N = 424) and the sole effect of Ts/Tv (Spearman's rho = 0.23, p-value = 2.021e-06; N = 424), suggesting that this second component could reflect a complex signature of a specific mutagen associated with generation length. Indeed, both transversions $T_H$>$A_H$ and $C_H$>$A_H$, negatively associated with generation length (equation I), have strong effects on the second principal component and, as expected, point to the opposite direction compared to $A_H$>$G_H$ (Figure 2C left panel). Analysis of a subset of species with many (more than 60) mutations, used to reconstruct the mutational spectrum, demonstrated quantitatively similar results (data not shown).

To investigate if the second principal component is affected by the total number of mutations, used to reconstruct the mutational spectrum, we performed several analyses. We demonstrated that the second principal component positively correlated with the number of mutations (Spearman's rho = 0.2247093, p = 2.964e-06). However, in the multiple linear model where the second principal component was a function of both generation length and the number of mutations we demonstrated that effect of generation length is stronger and more significant:

PCA2 ~ -0.34*GL + 0.16*(Number Of Mutations); *equation (Ib)*
*N = 424, $R^2$=0.109,*
*total p-value* = 1.121e-11*; p-value (GL)* = 7.07e-09*; p-value (Number Of Mutations)* = 0.006*;*
*presented coefficients are scaled.*
The first principal component, loaded mainly by the most common $C_H$>$T_H$ transition, appears to represent an unknown source of variation, which we were unable to associate with any life-history traits such as generation length, body temperature, body mass, metabolic rate or the number of mutations used to derive the mutational spectrum.

(3.6) ratio of the two most common transitions:
(script: *VertebratePolymorphisms.MutSpecComparisons.Analyses.Ecology.Mammals.R*):

Since low frequency genetic variants might occur due to DNA damage during sequencing (Chen et al. 2017, 2018; Stewart et al. 2018) we replicated our results taking into account only two the most common transitions: $A_H$>$G_H$ and $C_H$>$T_H$. Analysing the fraction of $A_H$>$G_H$ (derived as $A_H$>$G_H$ / ($A_H$>$G_H$ + C>T)), we still observed the positive correlation of this fraction with generation length (Spearman's rho = 0.21, p = 8.441e-06, n = 424).

Similarly with PGLS analyses in suppl mat 3.3 we repeated three models, analyzing an association of $A_H$>$G_H$ / ($A_H$>$G_H$ + C>T) with generation length. The final model 3C shows a significant positive effect of the generation length on $A_H$>$G_H$ / ($A_H$>$G_H$ + C>T) and the absence of the effect of the 'number of mutations'.

| ***model*** | ***variable*** | ***coefficients*** | ***p values*** |
|---|---|---|---|
| ***3A: $A_H$>$G_H$ / ($A_H$>$G_H$+ $C_H$>$T_H$) ~ log2(generation length)**; N=211, lambda [ ML] = 0.000* | *intercept* | *-0.0353585* | *0.6318952* |
| | *log2(generation length)* | *0.0239113* | *0.0005575* |

| | | | |
|---|---|---|---|
| ***3B: $A_H$>$G_H$ / ($A_H$>$G_H$+ $C_H$>$T_H$) ~ 0 + log2(generation length)**;* *N=211, lambda [ ML] = 0.000* | *log2(generation length)* | *0.0206577* | *< 2.2e-16* |
| ***3C: AH>GH / (AH>GH+ CH>TH) ~ 0 + log2(generation length) + log2(number of mutations)**;* *N=211, lambda [ ML] = 0.000* | *log2(generation length)* | *0.0225146* | *6.665e-12* |
| | *log2(number of mutations)* | *-0.0034229* | *0.5373* |

(3.7) nucleotide-content-independent mutational spectrum demonstrates a positive correlation between the generation length and $A_H$>$G_H$:
(script: *VertebratePolymorphisms.MutSpecComparisons.Analyses.Ecology.Mammals.R*):

The fraction of $A_H$>$G_H$ in each species-specific mutational spectra depends on both the number of observed $A_H$>$G_H$ substitutions within a species and the number of $A_H$ nucleotides in fourfold degenerate synonymous position of *MT-CYB* of a given species, used for normalization (Figure S2A, 2A). To demonstrate that the observed results (Figure S2B, 2B, 2C) were not solely driven by the variation in the frequency of ancestral nucleotides, we recalculated $A_H$>$G_H$ fraction for each species using only substitutions from $A_H$: $A_H$>$G_H$ / ($A_H$>$G_H$ + $A_H$>$C_H$ + $A_H$>$T_H$). Using this approach we don't take into account nucleotide content of different species. We observed a positive correlation between $A_H$>$G_H$ and the mammalian generation length (Spearman's rho = 0.164, p = 0.0007, N = 424), suggesting that $A_H$>$G_H$ is higher in long-lived mammals irrespectively of the nucleotide content.

Similarly with PGLS analyses in suppl mat 3.3 we repeated models, analyzing an association of $A_H$>$G_H$ / ($A_H$>$G_H$+ $A_H$>$T_H$ + $A_H$>$C_H$) with generation length. The final model 4B shows a significant positive effect of the generation length on $A_H$>$G_H$ / ($A_H$>$G_H$+ $A_H$>$T_H$ + $A_H$>$C_H$) and the absence of the effect of the 'number of mutations'.

| ***model*** | ***variable*** | ***coefficients*** | ***p values*** |
|---|---|---|---|
| ***4A: $A_H$>$G_H$ / ($A_H$>$G_H$+ $A_H$>$T_H$ + $A_H$>$C_H$) ~ log2(generation length)**;* *N=211, lambda [ ML] = 0.000* | *intercept* | *0.6214988* | *1.759e-12* |
| | *log2(generation length)* | *0.0208230* | *0.007154* |
| ***4B: $A_H$>$G_H$ / ($A_H$>$G_H$+ $A_H$>$T_H$ + $A_H$>$C_H$) ~ log2(generation length) + log2(number of mutations)**;* *N=211, lambda [ ML] = 0.000* | *intercept* | *0.6581961* | *1.655e-08* |
| | *log2(generation length)* | *0.0196741* | *0.01515* |
| | *log2(number of mutations)* | *-0.0041067* | *0.62634* |

**(4) The long-term mutational bias affects nucleotide composition of complete mammalian mitochondrial genomes**

(4.1) The mutational bias affects neutral nucleotide composition of complete mammalian mitochondrial genomes:

Effect of mutational spectrum on neutral nucleotide content: high $A_H$>$G_H$ in long-lived species leads to decrease in $A_H$ and increase in $G_H$. Consideration of phylogenetic inertia by means of phylogenetically independent contrasts demonstrated the same trend (fraction of $A_H$: spearman's rho = - 0.09, p = 0.025; fraction of $G_H$: spearman's rho = 0.09, p = 0.021). Stepwise backward multiple linear model confirmed the importance of only these two nucleotides: in the final model, after the removing of all non-significant variables, generation length (GL) depends negatively on $A_H$ and positively on $G_H$:

*log2(GL) = 11.06 - 0.11*(fraction of $A_H$) + 0.46*(fraction of $G_H$), p-value of the fraction of $A_H$ is 0.023, p-value of the fraction of $G_H$ is <2e-16, $R^2$ = 0.229, presented coefficients are scaled.* equation (ii)

***Table S3.*** *Correlations between generation time and nucleotide fractions in neutral sites*

| class | method | nucleotide | Spearman's Rho / coefficient in linear model | P value |
|---|---|---|---|---|
| Mammalia (N = 650) | pairwise spearman's correlation | $T_H$ | -0.27 | 3.635e-12 |
| | | $A_H$ | -0.31 | 1.287e-15 |
| | | $C_H$ | 0.18 | 3.665e-06 |
| | | $G_H$ | 0.47 | < 2.2e-16 |
| | results of the backward stepwise multiple linear model | intercept | 11.06 | < 2.2e-16 |
| | | $A_H$ | - 0.11 | 0.023 |
| | | $G_H$ | 0.46 | < 2.2e-16 |

4.2) Nucleotide gradient along mtDNA is more pronounced in long-lived species

GitHub:
https://www.google.com/url?q=https://github.com/polarsong/mtDNA_mutspectrum/blob/WholeGenomesBranch/Head/2Scripts/WholeGenomeAnalyses.AtgcAlongGenomes.NoOverlap.R&sa=D&ust=1608842051864000&usg=AOvVaw2j8Fn-HlZ4RQmEyS4nQH8g

We were interested if the slopes of changes in nucleotide content (Figure 3) differed between short- and long-lived species. We plotted nucleotide content at neutral sites of each of 13 genes and correlated it with gene location, ranked from the shortest (MT-CO1) to the longest time (MT-CYB) of being single-stranded (Figure S3). We observed that the frequency of $G_H$ increased along the gradient of time of being single stranded (Spearman's Rho >= 0.62, p-values <= 0.028). Given the fact that this analyses was limited by the low number of genes (13), we performed more sensitive analysis splitting the genome of each species into 50 windows containing 25 neutral (fourfold degenerate synonymous) nucleotides and using only genes located on the same strand of the major arc (all genes except ND1, ND2 and ND6). For each window we estimated the frequencies of four nucleotides and for each species we calculated the Spearman Rho coefficient and p-value, estimating the changes in a given metric along the genome (Figure S3). We can see that the frequency of $G_H$ indeed has the strongest bias towards positive rho values, and additionally we can see also, that the second strongest bias belongs to $A_H$, which decreased along the genome (Figure S3).

To test if some of these gradients are more pronounced in long- versus short-lived mammals, we estimated the fraction of long-lived mammals (with generation length more than median, 1101.1 day) among significant (with p-values <=0.01) rho values. Only for $G_H$ nucleotides we observed significant excess of long-lived mammals among the significant ones (Figure S3, odds ratio = 1.61, p-value = 0.003521 for $G_H$ nucleotide, Fisher test). This means that the rate of increase in the frequency of $G_H$ is faster in long- versus short- lived mammals.

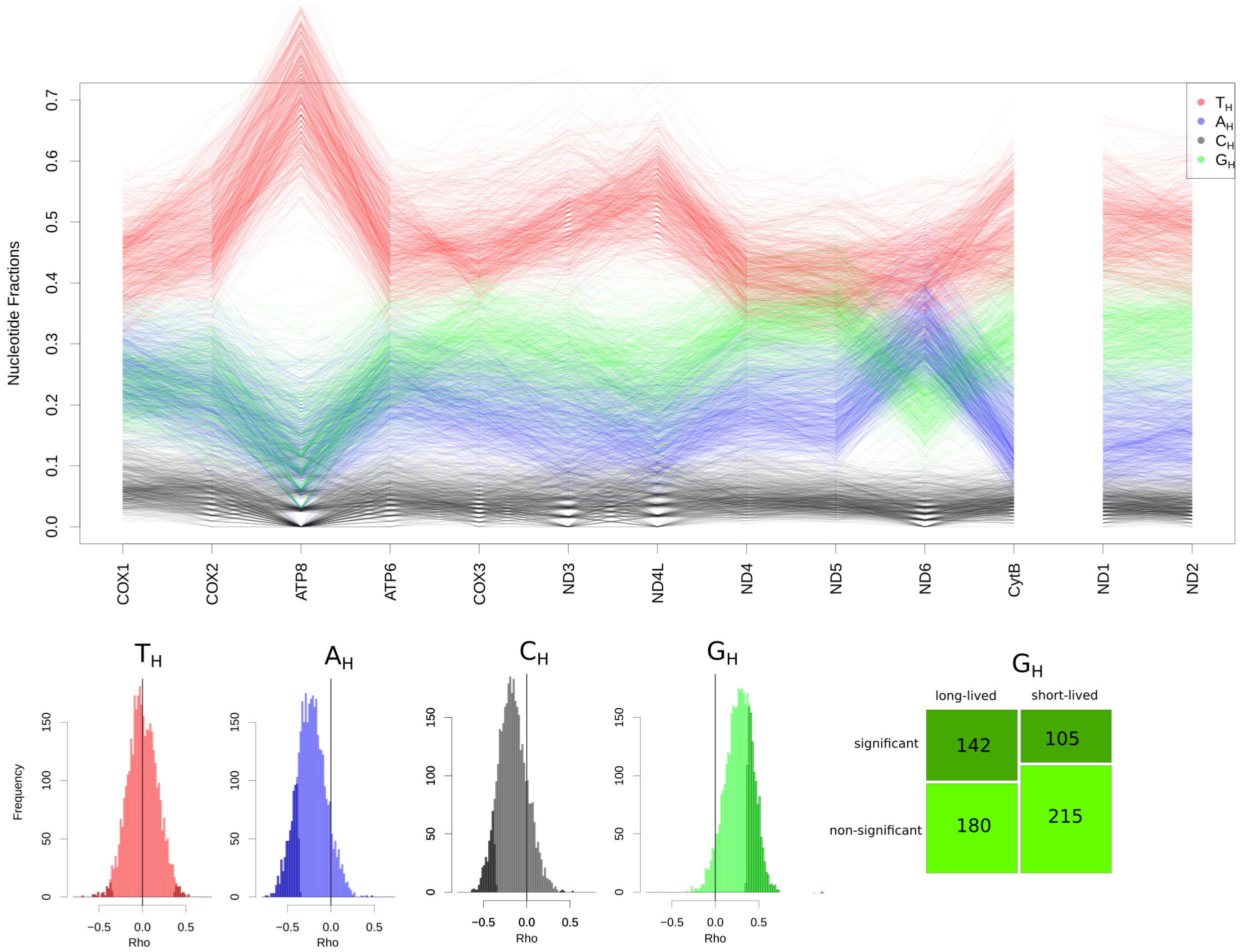


*Figure S3. The long-term effect of the mutational bias: neutral nucleotide content in mammalian species*

*Upper panel. Changes in nucleotide content along mammalian mtDNA. (N = 650). All genes located in the major arc are ranked according to the time spent single stranded: from COX1 to CYTB. ND2 gene spent more time than ND1 in the single-strand state but we do not compare directly these two genes with all others from the major arc.*

*Lower panel. The gradient of nucleotide changes with time being single stranded (increase in $G_H$) is more pronounced in long-lived mammals (N = 650). Histograms of spearman Rho values demonstrate that $A_H$ is decreasing with the time being single stranded while $G_H$ is increasing.*

*Mosaicplot reflects an excess of long-lived mammals among species with significant Rho for $G_H$, reflecting the higher* rate of increase in the frequency of $G_H$ *in mtDNA of long-lived mammals.*
